# Environmental heterogeneity plays a bigger role than diet quality in driving divergent California sea lion population trends

**DOI:** 10.1101/2025.04.23.650199

**Authors:** Ana Lucía Pozas-Franco, David. A. S. Rosen, Andrew W. Trites, Francisco J. García-Rodríguez, Claudia J. Hernández-Camacho

## Abstract

While the global population of California sea lions (Zalophus californianus) is increasing, regional trends show a decline in the Gulf of California (GoC, Mexico) and an increase in the Channel Islands (CI; California, U.S.) over the last 40 years. The drivers of these divergent trends remain unclear, but previous pinniped studies suggest that differences in diet quality—rather than prey abundance—may play a role. We therefore conducted an analysis to examine how sea lion population trajectories relate to diet quality, specifically looking at diet energy density and diet diversity. Using population and diet data from 1980 to 2020 for sea lions in the GoC and CI, we found no simple relationships between population trajectories and diet quality over time. Energy densities of sea lion diets were similar between the two regions, but GoC sea lions consumed a more diverse range of prey (n = 88 vs. 23 main prey species) dominated by benthic species and schooling fishes, while CI diets consisted mainly of schooling fishes and squid. We also found that GoC sea lions ate more benthic prey and less schooling fish during the 2014–2016 heatwave— decreasing their overall diet energy density, similar to the CI. This shift coincided with a temporary population decline in the CI but had variable effects on GoC populations. Overall, our findings suggest that regional population trends are influenced by complex ecological factors beyond diet quality alone, highlighting the need to consider environmental variability and prey composition when assessing the resilience of sea lion populations to climate-driven changes.

## 1. Introduction

California sea lions (*Zalophus californianus*) are widely distributed along the Pacific coast of North America from British Columbia, Canada to the Gulf of California, Mexico, but only reproduce on certain islands (rookeries) along the southern coast of California (U.S.), the Mexican Pacific, and the Gulf of California (1,2). Most of the global population (80%) breeds in California where numbers increased at an annual rate of 2.9% between 1964–2014 (3). The remainder of the population (20%) breeds in Mexico, where—with a few exceptions—most rookeries show a declining trend. In the Gulf of California, sea lion populations experienced on average a 2% decline per year between 1984–2015 (4,5).

In the southern Pacific coast of the U.S., California sea lions breed almost exclusively at four rookeries that form part of the Channel Islands which vary in size from 3,000–60,000 individuals (6). This population has grown steadily since the 1980’s but has experienced temporary population declines in some years associated with increased sea surface temperatures as seen during the 2012– 2016 marine heatwave (6). The population has since recovered, totaling 111,713 sea lions in 2019 (6). In contrast, breeding sea lion populations in the Gulf of California, Mexico, are distributed among 13 rookeries of varying size from 400–6,000 individuals (1,2). Although these rookeries vary in population growth, most show a declining trajectory since the 1980’s. Only one rookery in the southernmost Gulf of California, Los Islotes, is considered to have a population that has been increasing since 1979 (7).

The underlying factors causing a divergence in California sea lion population trajectories in the Channel Islands compared to the Gulf of California are still unknown (5). Possible contributing factors that are known to affect marine mammal species in general include regional differences in prey availability, pollution (both chemical and noise), disease, biotoxins, fishing gear entanglements, anthropogenic mortality (disturbance, legal and illegal shooting), and migration (8–11). Of these contributing factors, regional differences in diets associated with environmental change have been identified as the most likely contributor to population trajectories in several species (4).

Previous studies have generally focused on the negative effects that short-term reductions in prey *abundance* have on California sea lion numbers at rookeries in the Channel Islands (3,12,13) and the Gulf of California (7,14–18). Sharp declines in quantities of primary prey species available to sea lions are known to occur during El Niño events in California when warm water causes prey to remain at inaccessible depths, leading to increased pup mortality (19). However, El Niño events do not appear to have a comparable direct effect on the Gulf of California sea lion populations where the response to warming events depends on location, population size, and regional dynamics (5). This leads to the question of whether changes in the *quality* of prey (rather that changes in *quantity*) might better explain differences that have occurred in sea lion numbers over a longer timeframe (20), as suggested for Steller sea lions (21–25).

Diet quality can be assessed in many ways. Two main metrics of diet quality are diet energy density (an important aspect of the nutritional value of prey species) and diet diversity (i.e., the variety of species that compose the diet). These diet characteristics can affect the nutritional status of individuals, their reproduction and survival rates, as well as their susceptibility to disease and predation (23,24). Based on broad trends observed among different marine mammals, a diet dominated by a few energy-rich species would be hypothesized to support a growing population, while a switch to a more diverse diet of energy-poor species would be expected to cause population declines (26). It has also been demonstrated that changes in environmental conditions can alter the diet composition and ‘quality’ of prey species available to predators. However, it is not known how such differences or changes in diet quality may be influencing the population dynamics of California sea lions in the Channel Islands compared to the Gulf of California.

To investigate the effect of diet on population trajectories, we used estimates of average diet energy density, and two measures of diet diversity to quantify *diet quality* of California sea lions at the Channel Islands and the Gulf of California rookeries from 1980–2020. We compared the measures of diet quality between and within the two geographic regions and tested for relationships between rates of population change and the different measures of diet quality over time. Finally, the effects of increased sea surface temperatures (2014–2016) on sea lion diet and populations were also compared between regions. Obtaining a better understanding of the interplay between environmental changes, diets, and population trajectories is needed to inform policies regarding the conservation and management of California sea lions in Mexico and the U.S.

## 2. Materials and Methods

### 2.1 Population and diet data

This meta-analysis focused on the four California sea lion rookeries in the Channel Islands (San Miguel, San Clemente, Santa Barbara, and San Nicolas), and 12 of the 13 rookeries along the Gulf of California (Fig.1). We omitted San Jorge due to a lack of available data and also had insufficient data to include the west coast of the Baja California Peninsula. Available population and diet data for California sea lion rookeries in the Channel Islands and the Gulf of California from 1980–2020 were gathered from published and unpublished sources (Table S1). All data (population counts and diet data) used in this study were collected during the sea lion breeding season from May to August because: 1) most of the data available was from this season, 2) population data from this season captures the maximum number of sea lions present that year at a rookery by including newborn pups, and 3) we wanted to avoid the potential confounding effects that might be introduced by seasonal changes in diet when comparing diets across years and areas.

**Fig. 1.**
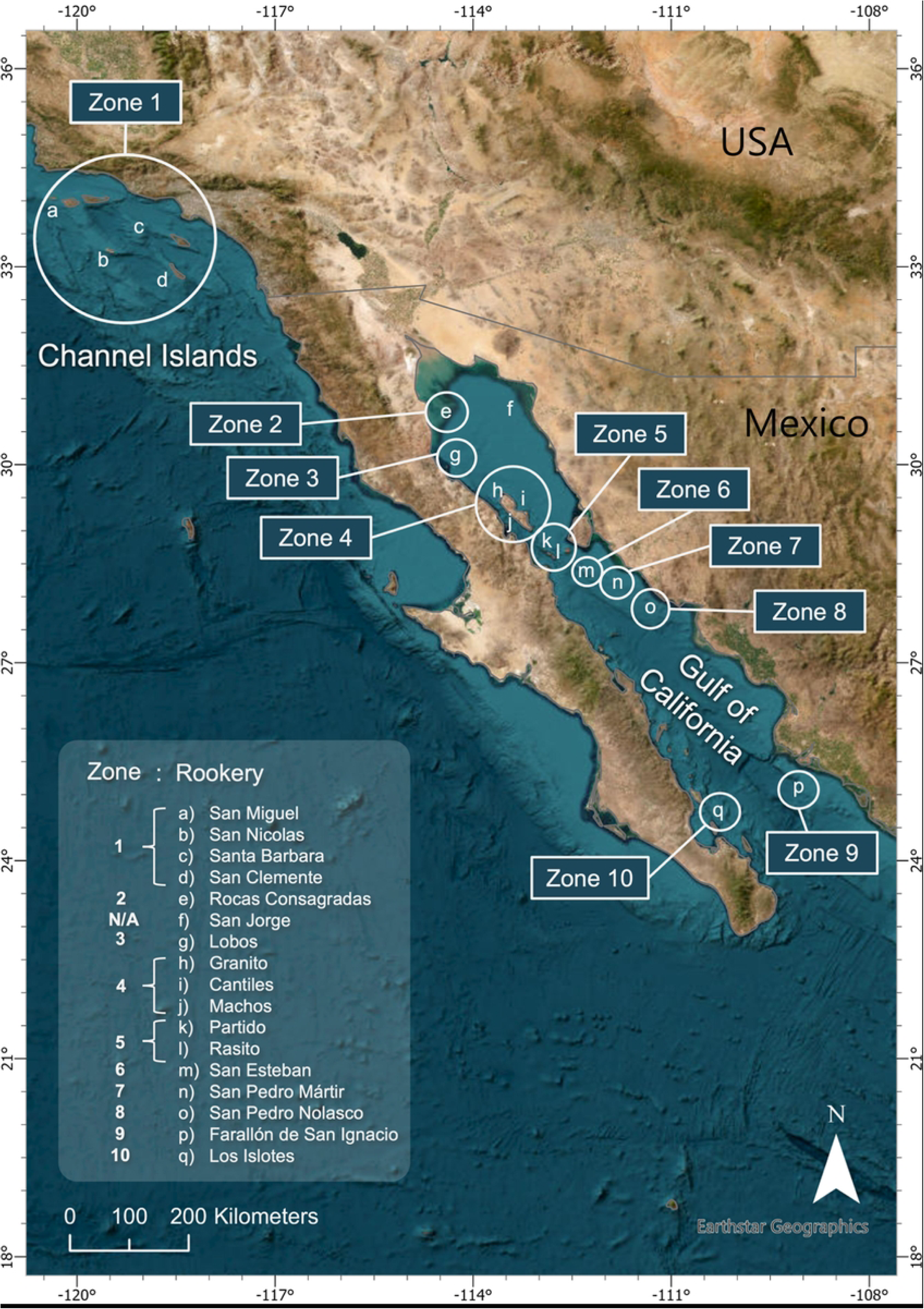
Map of the California sea lion rookeries and designated Zones for this study. Study sites included the four rookeries in the Channel Islands (a–d: San Miguel, San Nicolas, Santa Barbara, San Clemente) designated as Zone 1, and the 13 rookeries along the Gulf of California (e–q), and their respective Zones (2–10). Circles indicate rookeries within the indicated Zone. Rookery f (San Jorge) did not have diet data available and was omitted from further analysis. For reference, the distance between the northernmost rookery of Roca Consagradas (Zone 2), and the southernmost rookery of Los Islotes (Zone 10) is 823 km.

#### 2.1.1 Diet data

Diet data refers to data on the occurrence of identified prey species from sea lion scat samples collected at rookeries. Available data was used to assess diet quality, which we characterized using measures of diet diversity and diet energy density. Both diet characteristics were calculated using data originally reported as frequency of occurrence of prey species (FO) (27), or as an index of importance (IIMP) of prey species (Gulf of California data only) (16,28).

To test the relationship between diet quality and population change, we ideally needed matching diet data for the years with available population data. However, diet data were sparse, unevenly distributed over time, and only available for certain rookeries and years. Additionally, in some cases data were reported as a single mean spanning several years (e.g., Santa Barbara Island, 1981– 1995; Table S1). To address these limitations and make use of all available data, various data processing methods were employed prior to conducting analyses.

#### 2.1.2 Processing diet data

Only prey species with FO values ≥5% were included in this dataset because: 1) this was the cut-off available from most of the data, and 2) it served to highlight the main prey items. We also applied a ≥5% cut-off to IIMP values to maintain consistency across the data (this is the same IIMP cut-off previously used by Porras-Peters 2008). Note: while some literature uses a cut-off of IIMP values ≥10% (16), we were able to apply a ≥5% IIMP value cut-off because we had access to the raw diet data (F.J García-Rodriguez unpubl. data). ‘Non–identified’ species reported in the IIMP data were deemed not useful for this analysis and were therefore excluded.

To be able to compare diet data between rookeries and years, remaining FO and IIMP values were standardized to sum to 1.0 (or 100%) within each year of data by dividing each reported value by the total FO or IIMP for that rookery and year. This yielded *modified* frequency of occurrence (MFO; (29) and *modified* importance index (MIIMP) values which were used from this point onwards to calculate diet diversity and energy density.

#### 2.1.3 Calculating diet diversity

Two measures of diet diversity were used: 1) the total number of species recorded in the diet, and 2) diet diversity calculated using the Shannon Index (30) from either MFO or MIIMP values for individual species, where a higher resulting H-index indicates greater species diversity. Values of average diet diversity using the Shannon Index were calculated for each rookery and year when data was available, which were then used in subsequent analyses (Table S2).

#### 2.1.4 Calculating (weighted) diet energy density

The energy density of each prey species was recorded in kilojoules per gram of wet weight (kJ/gww) from data obtained in Gleiber et al., 2022 (31). If an energy density value was not available at the species level, an average energy density value from the species’ family (or in a few cases a closely related family) was used instead to approximate energy density. Then we calculated the *weighted* average diet energy density (to account for relative appearance of each prey species in the diet) by multiplying the MFO or MIIMP (expressions of proportion in the diet) by the respective energy density of that prey species (in kJ/gww). Summing these values gave an average weighted diet energy density value for each rookery and year, which was used in subsequent analyses (Table S2).

#### 2.1.5 Population data

All sea lion counts from 1980–2018 from the Gulf of California rookeries were obtained from (17). Counts were made from boat surveys and included numbers for each age and sex class. Population counts for the Channel Islands were available from 1980–2019 and were sourced from (3) for 1980–2014, from (32) for 2015, and from (6) for 2016–2019 (Table S1). These counts had already been corrected for pups that were obscured from vision and for adult females that were foraging during the census. At times, multiple sources reported population counts for the Channel Islands, so the source with the higher counts was used if this was because a technique with presumed greater accuracy was used (i.e., aerial photography counts vs. boat counts).

#### 2.1.6 Calculating population change

To test the relationship between population and diet quality, we calculated rates of population change for years with available diet data at each rookery. Using regression analysis, we estimated population changes immediately associated with single or grouped years with diet data (i.e., year-rookery grouping) by incorporating population counts from a set number of years before and after the diet data. Since the range of years incorporated into each rookery-year diet data point varied, a set of rules were established to define the number of years before and after the diet data span that were included in the calculation of population change, depending on how many years of diet data were included for that grouping. Further details on calculating population change provided in the Supplementary Methods. In general, population trends were estimated from data that spanned the period of diet data by at least an additional year on either side.

### 2.2. Grouping data

#### 2.2.1 Creating Zones and Zone-era groupings

To investigate the relationship between population changes and diet quality, we had to ensure the independence of the data points on both a temporal (both sequential and non-sequential data) and geographic scale (closely related rookeries). This involved grouping rookery diet data (energy density and diet diversity averages) and population data (population change averages over a set of years) together into non-continuous sets of years into rookery-year groupings (more details in Supplementary Methods). Then we created sets of matching population and diet data sets averaged across related geographic areas (Zones) and non-continuous time periods (eras): Zone-era groupings.

As previously noted, neither population or diet data was continuous across years, and the timing of the data was not consistent across rookeries, resulting in the previously described rookery-year groupings. Values for the eventual ‘Zone-era’ data groupings were created by averaging the population change of each rookery-year grouping, and the respective diet quality values (energy density and diversity) within a geographic Zone (described below) for each set of number of years or ‘era’ with available data.

This grouping of data from individual rookeries into composite geographic Zones was done for two main reasons: 1) to prevent potential over-representation of individual rookeries relatively close to each other that could be considered common ecological units, and 2) to best deal with the lack of available continuous population and diet data over time by grouping available data, thus allowing us to make comparisons between different geographic Zones for similar time periods/eras. Rookeries were grouped into a common Zone if they occurred in a similar geographic area (<100 km away from each other) and had a similar population trajectory over time, which were determined by fitting a linear regression to the total population counts of each rookery for all years with available data (Fig. 2). The average annual population change (percent) was calculated as the slope divided by the intercept (the predicted first year population) of the regression equation. Population trajectories were therefore classified as increasing, decreasing, or inconclusive according to the slope of the linear regression.

**Fig. 2.**
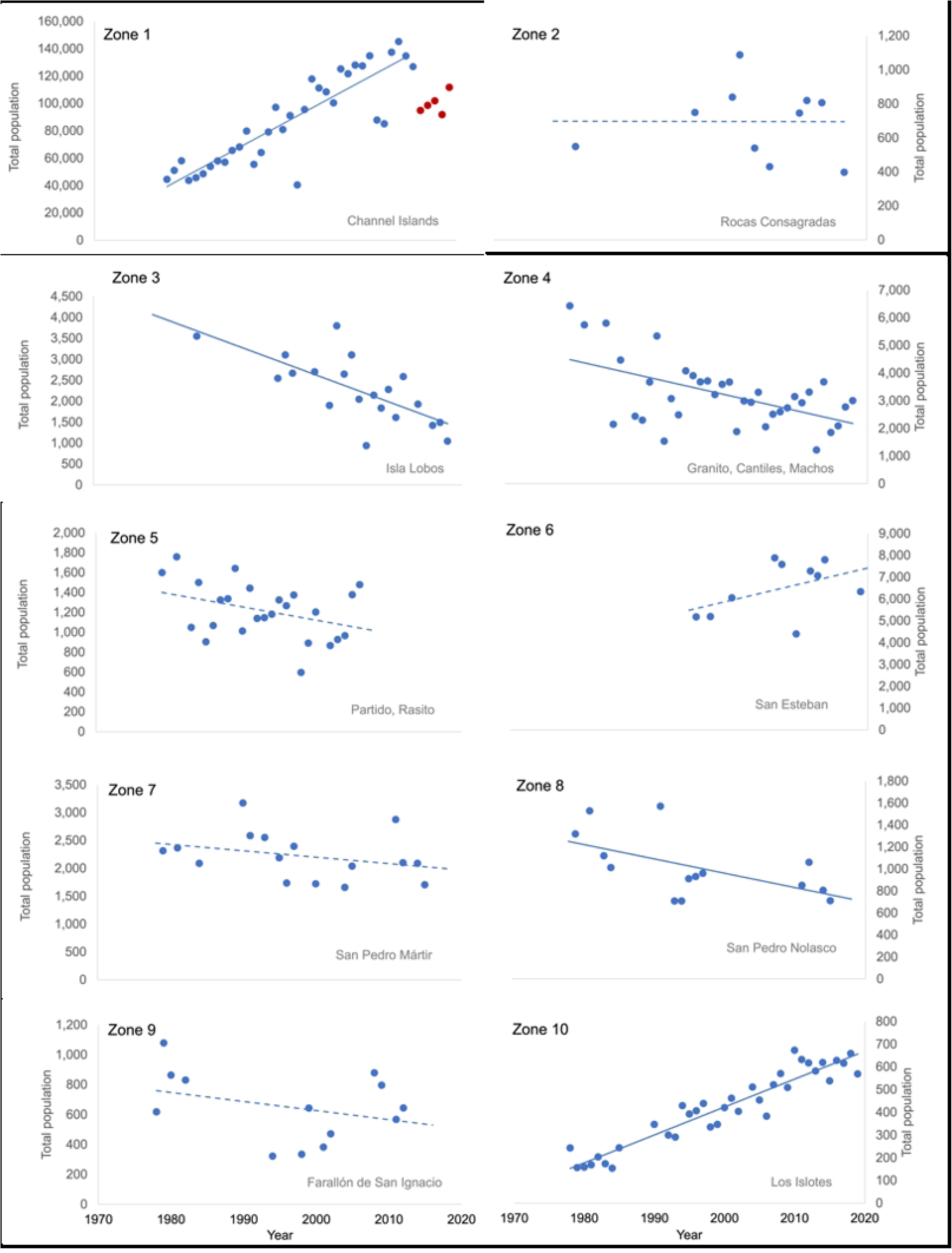
California sea lion population trends within Zones. (**1980–2020**). Data shows total sea lion counts for each Zone with data from 1980–2020. Rookery names within each Zone are labeled in the bottom right of each panel. Zones composed of multiple rookeries show the sum of the population totals in those rookeries. Solid lines represent statistically significant regressions and dotted lines represent regressions that were not statistically significant. Red data points in Zone 1 represent population declines after the 2014 warming event (2015–2019); these data were not included in the overall regression.

Previous studies have partitioned the 13 Gulf of California rookeries into three sub-populations based on factors such as environmental conditions, genetic structure, and diet (17,33–36). However, in our study we deliberately excluded factors related to diet, and thus created Zones based only on similarities in population trajectory and geographic proximity.

Following this methodology, 10 Zones were created: the Channel Island rookeries were grouped together into Zone 1 (Fig. 1). The available data resulted in two Zone-era groupings: Zone 1 1981– 1995 and Zone 1 2000–2011. The Gulf of California rookeries were grouped into 9 zones (Zones 2–10; Fig. 1). Data for the Gulf of California using the available FO data resulted in two era groupings, 1990–2000 and 2015–2019. This resulted in 9 Zone-era groupings for the GoC using FO data. The IIMP data yielded three era groupings: 1995–1996, 2002, and 2015–2019 (Table S3). This resulted in 18 Zone-era groupings for GoC. Hence, this process yielded 11 Zone-era sets of matched population and FO-based diet quality data for the CI and GoC combined, and an additional 18 sets of data for the Gulf of California from IIMP-based diet quality. The relationship between diet quality and population change was then tested on these ‘Zone-era’ groupings.

### 2.3 Effects of environmental change

Changes in diet quality before and after an event characterized by increased sea surface temperatures were explored by comparing the change in average diet energy density and diet diversity in the Gulf of California before and after 2014. Sufficient post-environmental shift diet data were not available for the Channel Islands post-environmental shift, so only pre-environmental shift data was used to compare the diet quality between the Gulf of California and the Channel Islands.

Prey species from the diet data were grouped into 9 species categories to further describe changes in diet before and after the environmental shift, and to express the ecological distribution of species consumed (number of species per category). These categories were assigned based on broad ecological characteristics similar to previous studies as per Trites et al., 2007 (37), and included: benthic species (n = 60 prey species), crustaceans (n = 1), gadids (n = 5), lanternfish (n = 7), octopus (n = 2), rockfish (n = 5), schooling fishes (n = 21), squids (n = 15), and miscellaneous (n = 17) (Table S4).

### 2.4 Statistical analysis

To test relationships between population changes and diet quality, linear regression models were fit to the data in R-Studio (version 2022.02.3). Simple linear models were used to test for relationships between population change and measures of diet diversity and energy density at the level of ecological Zones using Zone-era data. Additionally, since Zones consisted of varying population sizes, those regression analyses incorporated median Zone population size as a weighting factor. The resulting p-values and adjusted R-squared values from all linear models are reported. All energy density and diet diversity values derived from MFO and MIIMP data were tested for outliers using Grubb’s and Dixon’s outlier tests in R-Studio using the package “outliers”, at the Zone-era grouping level. Preliminary analyses revealed no statistically significant outliers in the data.

Two-sample t-tests assuming unequal variances were conducted using MFO and MIIMP data to compare diet diversity and energy density within the Gulf of California before and after the 2014 environmental shift, and when comparing pre-environmental shift diets between the Gulf of California and the Channel Islands.

## 3. Results

### 3.1 Population trajectories

All four Channel Island rookeries (Zone 1) averaged population increases of 2-6% per year from 1980–2020, except for a population decrease associated with increased sea surface temperatures in 2014 (Fig. 2). In contrast, the overall population in the Gulf of California decreased over the same time-period, although trends differed at individual rookeries. While most rookery populations showed a decline, San Esteban and Los Islotes rookeries (Zones 6 and 10) showed increasing population trajectories (although the San Esteban population increase was not statistically significant). As opposed to the Channel Islands, sea lion populations in the Gulf of California did not seem to be collectively affected by the 2014 warming event (Fig. 2).

### 3.2 Diet quality and population changes

Diet diversity (using the Shannon Index) did not correlate significantly with rate of population change when examined at the level of Zone-eras (*p* = 0.430 using MFO data and 0.622, using IIMP data; S3 Fig., left panels). Nor were there significant relationships between diet energy densities and rates of population change (*p* = 0.804 using MFO data and *p* = 0.128 using IIMP data; S3 Fig., right panels).

### 3.3 Diet quality in the Channel Islands vs. the Gulf of California

At the Channel Islands, California sea lions primarily consumed 23 main prey species, whereas in the Gulf of California, they consumed 88 main species (using MFO >5% data; Figs. 3 and 4). Jack mackerel, Chub mackerel, Pacific hake, and Californian anchovy coincided as top species consumed by sea lions in both the Channel Islands and the Gulf of California.

**Fig. 3.**
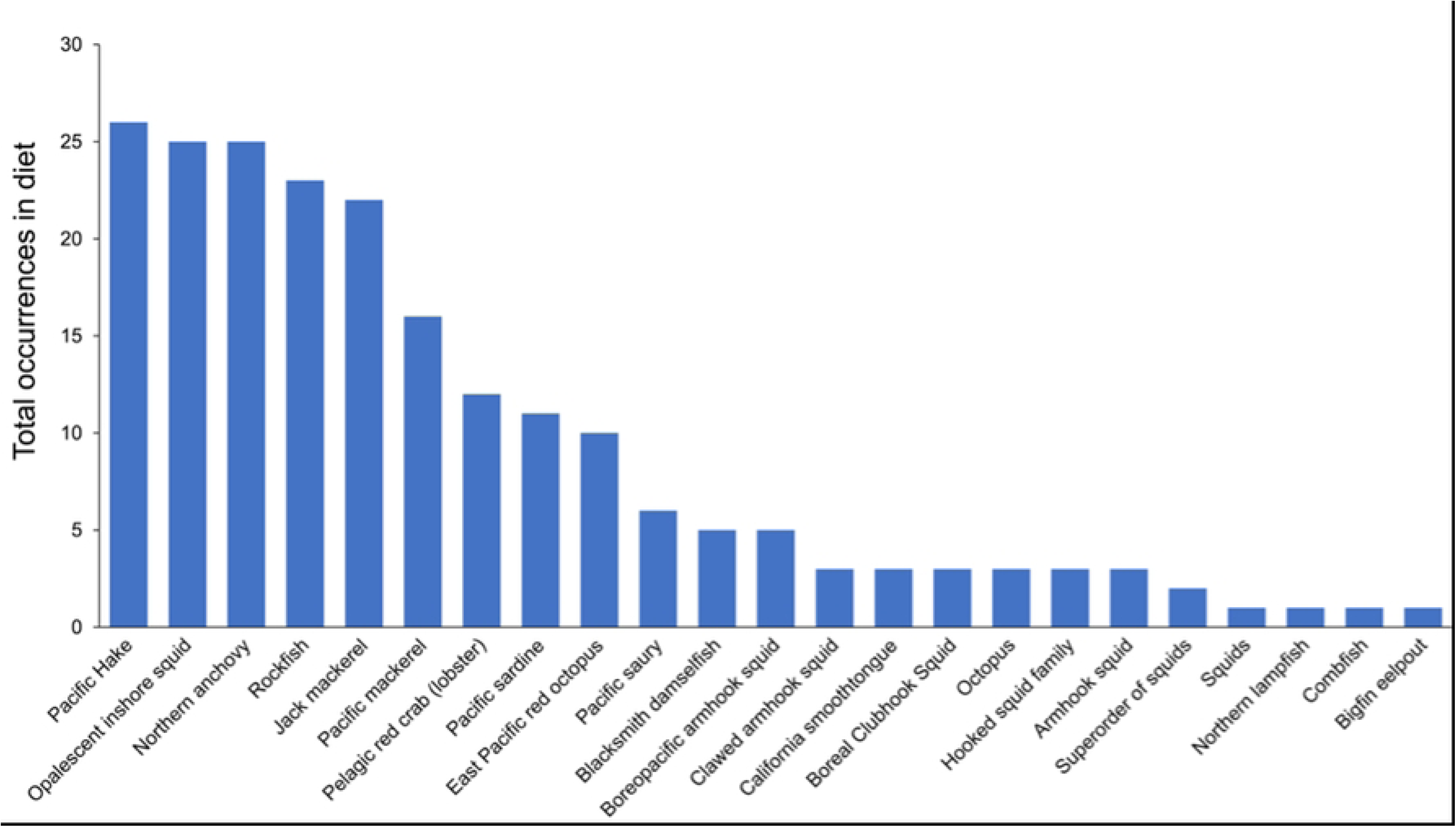
Prevalence of California sea lion prey species in the Channel Islands. Bars represent the total number of occurrences (out of 41 possible occurrences, equivalent to years) of each prey species from frequency of occurrence data from 1980–2011; that is, the total number of years where each prey species was present in the diet. All 23 species with FO ≥5% are listed. The species’ common name is listed when available, although some were identified only at the family or genus level.

**Fig. 4.**
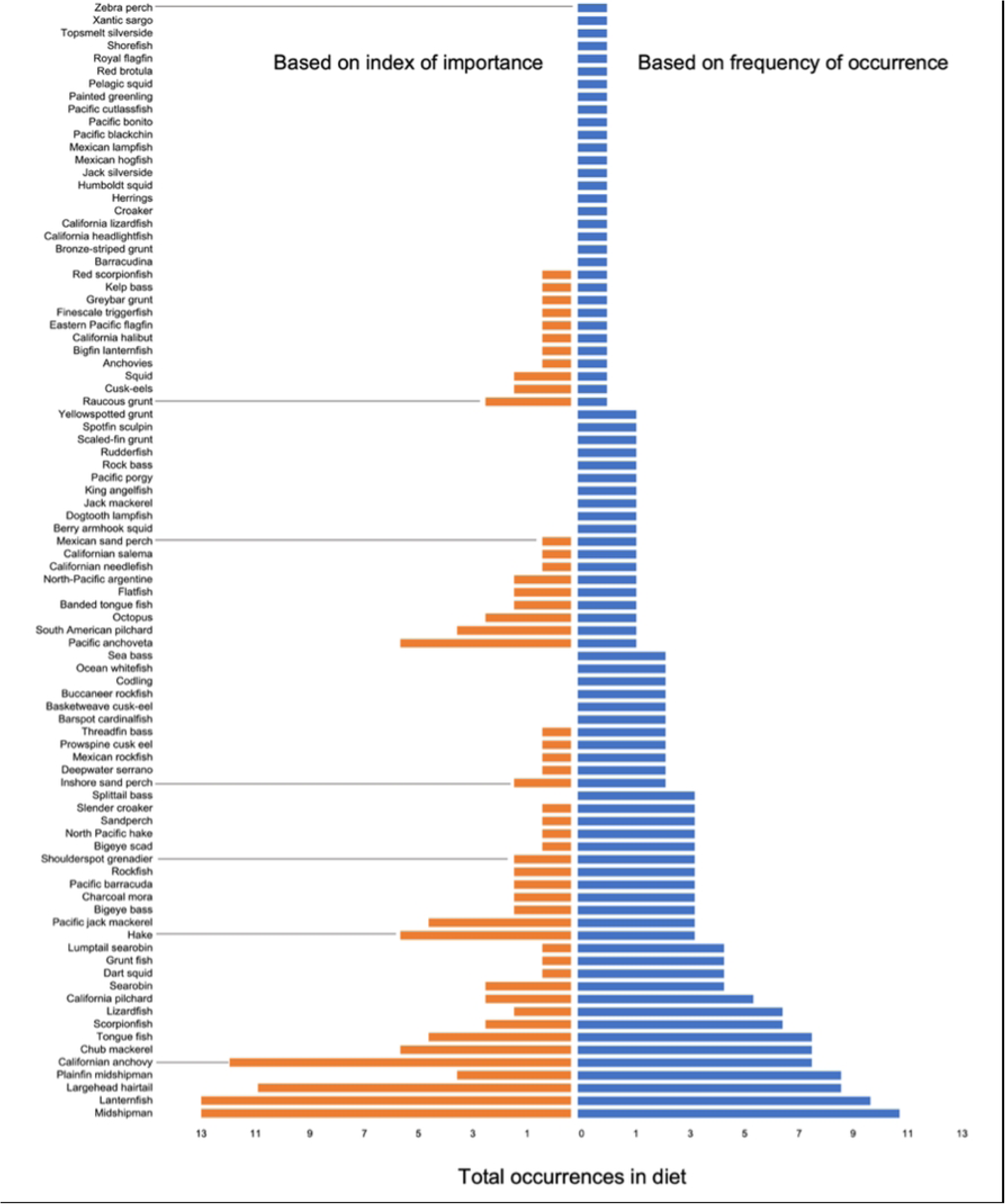
Prevalence of California sea lion prey species in the Gulf of California. Bars represent total prey species occurrences in the diet per year (total number of years where each prey species was present in the diet), from available diet data from 1980–2019 for all Gulf of California rookeries (Zones 2–10) from either frequency of occurrence (blue bars) or index of importance (orange bars). The common name of all 88 species (or species groups) from both MFO and MIIMP data is listed when available (some species were identified only at the family or genus level).

Sea lions in the Channel Islands ate mainly schooling fishes and squid (36% and 21% of diet respectively; data from 1981–2011), whereas sea lions in the Gulf of California mainly ate multiple benthic species (41 species; 47% of diet) and schooling fishes (19 species; 22% of diet; data from 1990–2019; Fig. 5, left panels), with schooling fishes having a higher average energy density than benthic species (Fig. 6).

**Fig. 5.**
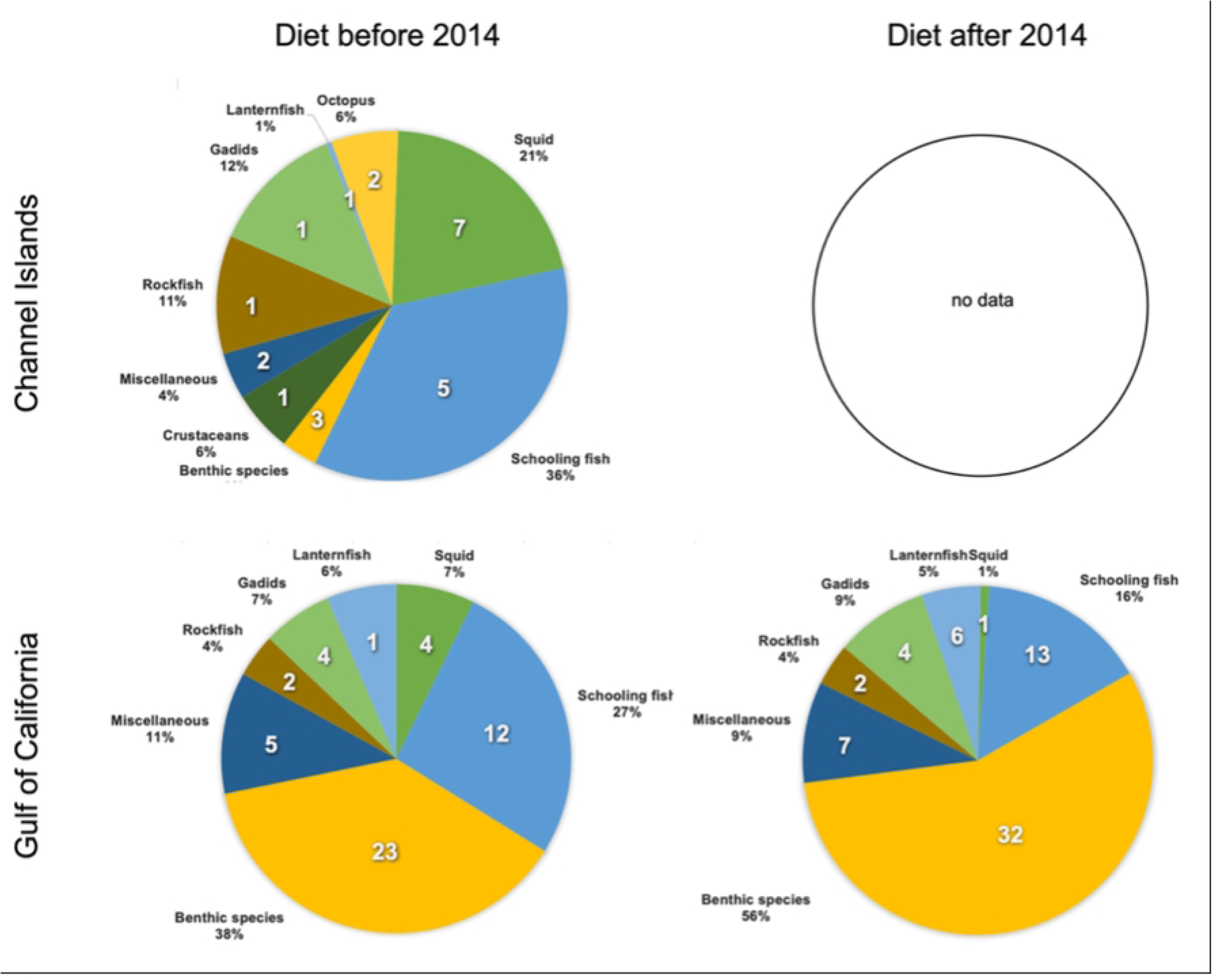
Diet composition by prey species categories before and after 2014 from frequency of occurrence data. Pie chart slices represent the proportion by each species category. White numbers represent the number of species in the diet from each category. Diet composition data from the Channel Islands is from 1980–2011 (no comparable diet data available after 2011). Diet composition data for the Gulf of California before the 2014 environmental shift is from 1990–2000 and from 2015–2019 after 2014.

**Fig. 6.**
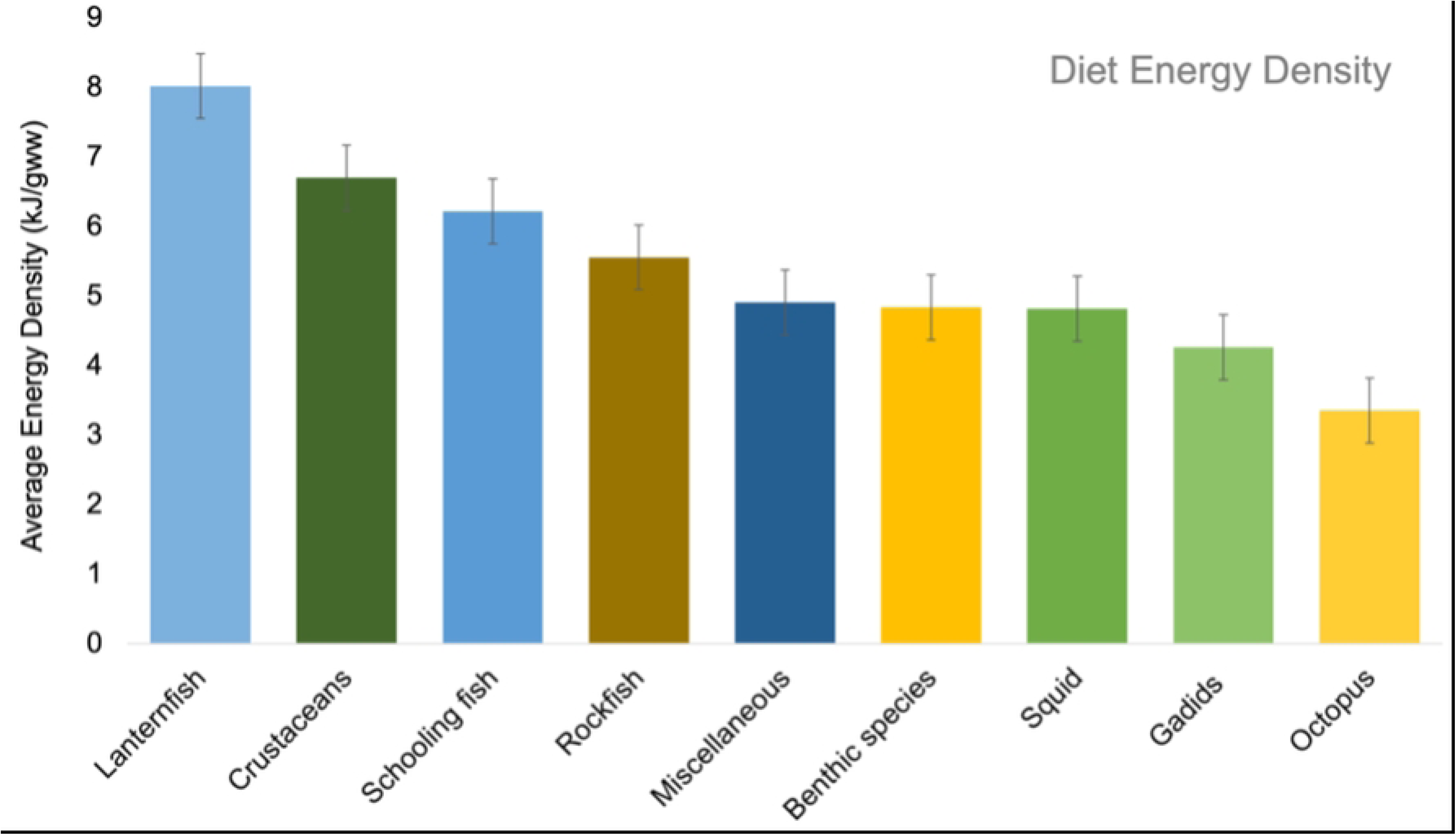
Average energy density of each prey species category consumed by California sea lions. Colours correspond to species categories illustrated in Fig. 6. Bars represent average energy densities (kJ/gww; mean value ± standard error) from all species present in the diet data from each category ordered from highest (lanternfish) to lowest (octopus) value.

Mean diet energy density for sea lions in the Channel Islands (5.43kJ/gww) was comparable to the Gulf of California (5.32 kJ/gww) (Table 1 and Fig. 7). Mean diet diversity using the Shannon Index was lower in the Channel Islands than in the Gulf of California (1.83 vs. 2.04; Table 1 and Fig. 7) and had higher variability among the Gulf of California Zones (range: 0.79–3.26) than among the Channel Islands (range: 1.34–2.35).

**Fig. 7.**
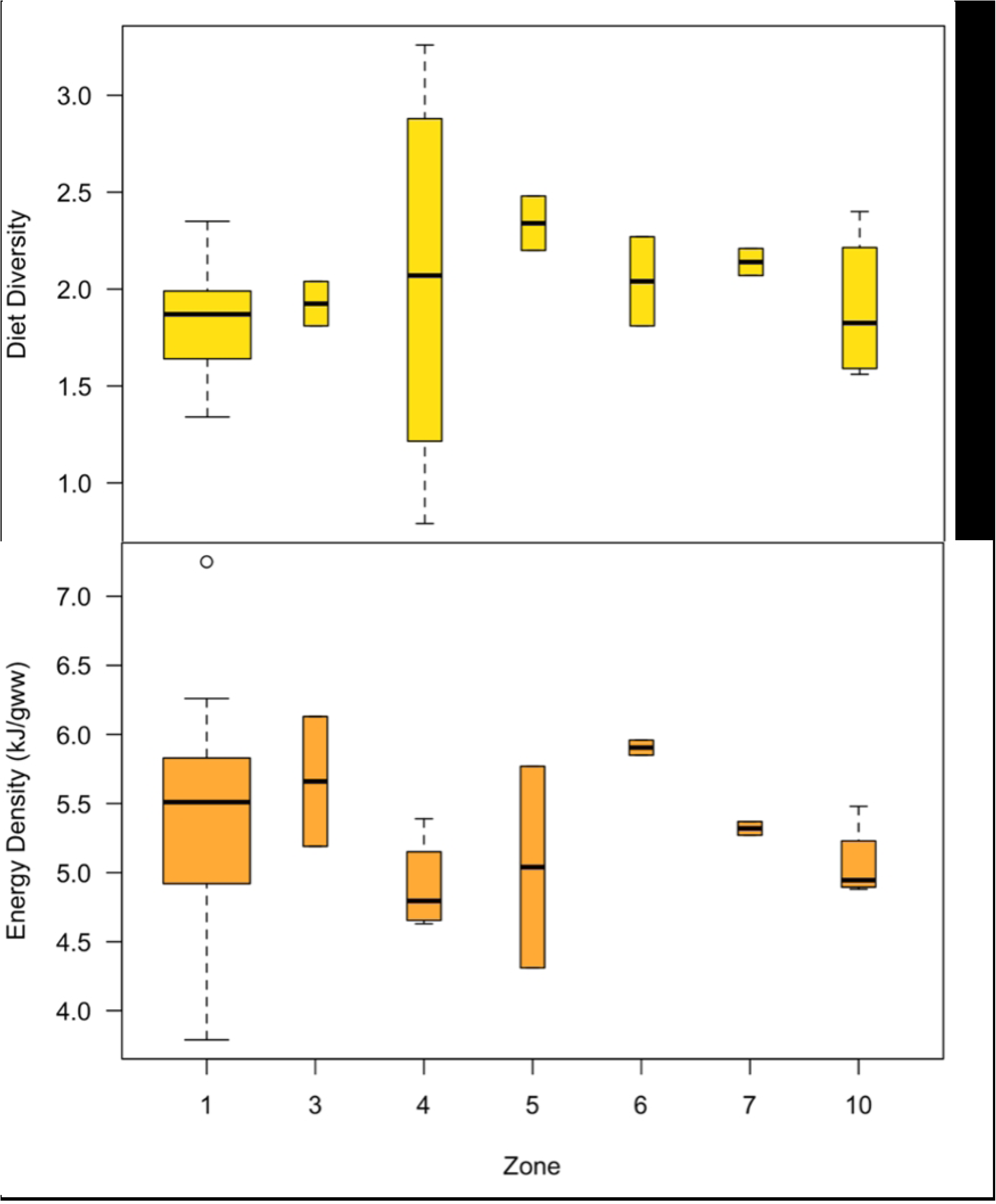
Average annual diet diversity and energy density by Zones from frequency of occurrence data. Diet diversity (top panel) and energy density (bottom panel) values were based on all available Zone and year groupings from MFO (1981–2019; Zones 2, 8 & 9 are omitted due to lack of available FO data). Diet diversity values were calculated using the Shannon Index. Box limits represent the first, mean, and third quantile values, box whiskers represent the range of values. Bar widths are proportional to the number of data points in each Zone. The circle represents the energy density value of 7.25 in Zone 1 (San Miguel, 2005) which was considered an outlier according to Grubb’s test.

**Table 1.** Diet quality and population trajectory of each of the Gulf of California Zones and the Channel Islands from frequency of occurrence data. The mean and standard deviation of the range of the annual weighted diet energy density (kJ/gww) and of the diet diversity (Shannon Index) for sea lion diets is shown (incorporating individual prey species). ‘n’ represents the number of years with data points used to calculate the corresponding mean and range values. Population trajectories in **bold** represent statistical significance. Zones 2, 8 & 9 are omitted due to lack of available FO data.

| Zone: rookery | Population trajectory | Energy density (kJ/gww) |  | Diet diversity (Shannon Index) |  | n |
| --- | --- | --- | --- | --- | --- | --- |
|  |  | Mean ±SD | Range | Mean ±SD | Range |  |
| 1: San Miguel | Increasing | 5.43 ±0.94 | 4.14–7.25 | 1.79 ±0.94 | 1.34–2.35 | 10 |
| 1: San Nicolas | Increasing | 5.63 ±0.22 | 5.26–5.94 | 1.75 ±0.23 | 1.46–1.96 | 8 |
| 1: San Clemente | Increasing | 5.16 ±0.84 | 3.79–6.05 | 1.94 ±0.84 | 1.67–2.21 | 7 |
| 1: Santa Barbara | Increasing | 5.48 | N/A | 1.87 | N/A | 1 |
| **Channel Islands average** | **Increasing** | **5.43 ±0.20** | **3.79–7.27** | **1.83 ±0.12** | **1.34–2.35** | **4** |
| 3: Isla Lobos | **Decreasing** | 5.66 ±0.66 | 5.19–6.13 | 1.93 ±0.66 | 1.81–2.04 | 2 |
| 4: Machos, Cantiles, Granito | **Decreasing** | 4.90 ±0.35 | 4.63–5.39 | 2.05 ±0.35 | 0.79–3.26 | 4 |
| 5: Rasito | Decreasing | 5.04 ±2.45 | 4.31–5.77 | 2.34 ±1.03 | 2.20–2.48 | 2 |
| 6: San Esteban | Increasing | 5.90 ±2.83 | 5.85–5.96 | 2.04 ±0.08 | 1.81–2.27 | 2 |
| 7: San Pedro Mártir | Decreasing | 5.32 ±2.50 | 5.27–5.37 | 2.14 ±0.07 | 2.07–2.21 | 2 |
| 10: Los Islotes | **Increasing** | 5.06 ±2.17 | 4.88–5.48 | 1.90 ±0.28 | 1.56–2.40 | 4 |
| **Gulf of California average** | Decreasing | **5.32 ±0.39** | **4.31–6.13** | **2.04 ±0.15** | **0.79–3.26** | **6** |

The lowest diet diversity (0.79) occurred within Zone 4 (Granito, Cantiles, Machos) in 1996, which was heavily influenced by the Granito rookery where just one main species, largehead hairtail (*Trichiurus lepturus*), was consumed that year. Interestingly, the highest diversity recorded among all locations (3.26) occurred in 2018 also within Zone 4 where data were exclusively from Granito and showed that that year, sea lions consumed a total of 31 main species.

The mean diet energy density in the Channel Islands was 5.43 ±0.2 kJ/gww, with a surprisingly small overall variation considering that two data points had anomalously high or low energy densities (7.25 kJ/gww in San Miguel, 2005 and 3.79 kJ/gww in San Clemente, 1982). The species that contributed the most to the average energy density in the diet primarily belonged to the schooling fishes category (Fig. 6), and included jack mackerel, Pacific mackerel, and northern anchovy. San Miguel Island in particular had years when sea lion diets had above-average energy densities (S1 Fig.) with a higher-than-normal contribution from two species of schooling fishes, Pacific sardine (2002–2005) and herring (2005).

Although the overall diet energy density for sea lions in the Gulf of California was not statistically different than for those in the Channel Islands, there was significant variability between years and within Zones in the Gulf of California, demonstrating differences in the species that contributed the most to the average diet energy densities. The highest diet energy density in the Gulf of California in any year occurred in Zone 3 (Isla Lobos, 1995, 6.13 kJ/gww), mainly due to the high energy density of Pacific anchoveta (a schooling fish) and largehead hairtail (a miscellaneous fish) (Table 1 and S2 Fig.). However, diets in Zone 6 (San Esteban) had the highest mean energy density overall (5.90 kJ/gww; 1995 and 1996). For both Zones 6 and 7 (San Pedro Mártir), the overall energy density of the diet largely reflected a high contribution from lanternfish, followed by largehead hairtail, Californian anchovy, and chub mackerel (S2 Fig.). Interestingly, in 1996 lanternfish was largely replaced with other species in the diet in both Zones 6 and 7 with little effect on the mean diet energy density. Diets of sea lions feeding in Zone 4 (Granito, Cantiles, Machos) had the lowest mean energy density (4.90 kJ/gww), which included a high proportion of ‘other’ species outside of the top 17 for most years (1996, 2016, 2018) (S2 Fig.).

### 3.4 Effect of environmental changes on diet quality in the Gulf of California

From 2014 to 2016, the Gulf of California experienced unusually high sea surface temperatures. During this period, the proportions of schooling fish and squid in the diet of California sea lions decreased significantly, from 27% to 16% and from 7% to 1%, respectively, while the proportion of benthic species increased from 36% to 56% (Fig. 5). These changes marked a shift from high-energy schooling fish to lower-energy-density benthic species, resulting in a significant reduction in the overall average energy density of their diet to 4.69 ± 0.35 kJ/gww (two-tailed p = 0.045; Table 2, Fig. 8, bottom panel).

**Fig. 8.**
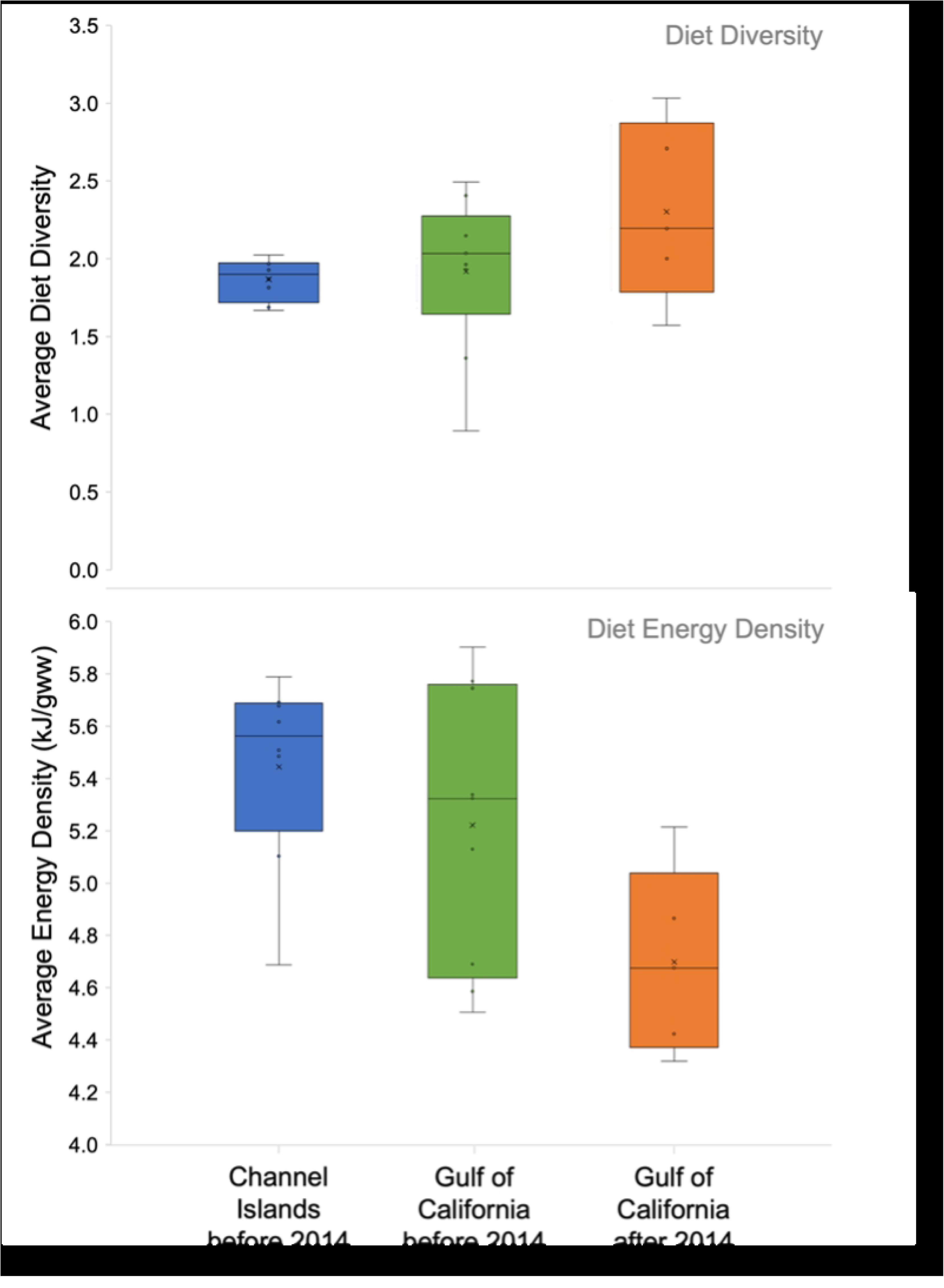
Average diet diversity from the Shannon Index (top panel) and average energy density (bottom panel) from frequency of occurrence data before and after 2014. Box limits represent averaged data (first, median and third quantiles ± standard error, ‘x’ represents mean value) from all Channel Island and Gulf of California groupings before and after 2014 (no data for Channel Islands after 2014). Diet data is based on rookery-year groupings. No significant differences were found between mean diversity values between geographic areas before 2014, nor in the Gulf of California between eras. There was a statistically significant decrease in the average energy density in the Gulf of California after 2014.

**Table 2.** Diet quality by geographic area before and after 2014. Total number of diet prey species, mean diet diversity using the Shannon Index, and mean weighted diet energy density.

| Geographic area & era | Number of prey species in diet | Diet diversity (Shannon Index) (mean ±SD) | Energy density (kJ/gww; mean ±SD) |
| --- | --- | --- | --- |
| Channel Islands before 2014 | 23 | $1.83 \pm 0.12$ | $5.42 \pm 0.36$ |
| Gulf of California before 2014 | 51 | $1.92 \pm 0.49$ | $5.22 \pm 0.53$ |
| Gulf of California after 2014 | 65 | $2.30 \pm 0.57$ | $4.69 \pm 0.35$ |

Diet diversity calculated using the Shannon Index showed no significant difference in mean values before (1.92 ±0.49) and after 2014 in the Gulf of California (2.30 ±0.57; two-tailed *p* = 0.245; Table 2, Fig. 8, top panel). However, there was an increase in the number of prey species consumed (from 51 to 65 species), and an overall increase in the average number of prey species consumed per rookery after 2014 (S4 Fig.). Before 2014, sea lions consumed 9 prey species on average within rookeries (range 5–15 species), increasing to an average of 16 species per rookery (range 7–26 species) after 2014 (S5 Fig.; data unavailable for Zones 8 and 9). Equally notable was that ∼50% of the species consumed throughout the Gulf of California after 2014 were not present in the diet prior to this time (S6 Fig.) ––in other words, the sea lions did not simply add 14 more species to their diet but made a fundamental shift in the species they consumed. Despite the overall dietary shifts in prey quality observed after 2014, there were no apparent differences in the rates of population changes between these two eras (two tailed *p* = 0.984).

## 4. Discussion

Previous dietary studies on California sea lions in the Gulf of California have tended to focus on detailing differences in the main prey species consumed, or describing feeding behaviours at various rookeries (16,18). Only one study has indirectly explored the broad relationship between population changes and diet within the Gulf of California, finding no significant relationships between these variables (17). Our study is the first to assess California sea lion diet quality using available long-term summer data on a finer geographic scale for the Gulf of California (1990–2019) compared to the Channel Islands (1980–2014), to explore its relationship to differences in population trajectories.

Our results demonstrate substantial differences in the diversity and type of prey species consumed by sea lions in these two areas, but do not show any significant relationships between measures of diet quality (diet energy density or diet diversity) and long-term rates of population change. Results also demonstrate that following the significant increase in sea surface temperatures that occurred in 2014, sea lions in central regions of the Gulf of California consumed a greater number of prey species that had an overall lower diet energy density. However, this shift did not result in an overall population decline throughout the Gulf, as occurred in the Channel Islands. These findings underscore the importance of considering the environmental heterogeneity at the different regions throughout the Gulf of California, which can heavily influence California sea lion population dynamics at local levels.

### 4.1 The role of diet diversity

An ideal diet for California sea lions is one that allows them to meet their nutritional needs to grow and reproduce by feeding on sufficiently available prey species. However, our results illustrate that the exact nature of such a diet appears to vary depending on the characteristics of the ecosystem, making single indicators such as diet diversity difficult to interpret. Animals may choose to forage on fewer, energy-rich prey species (a low diversity, high energy density diet), which may reflect either a high degree of prey selectivity or a less biodiverse ecosystem. In more diverse ecosystems, foraging on a greater combination of species of different sizes and nutritional profiles might prove to be an optimal strategy and may even be required to fulfill nutritional requirements (38). Different marine mammal species have exhibited switches to lower quality diets (lower energy density prey) during environmental challenges that have been characterized by either decreases (22,39) or increases (40,41,26) in diet diversity.

Using the total number of prey species as a measure of diet diversity in our study revealed a striking difference in diets between regions. The sea lions at the Channel Islands consistently consumed 23 primary prey species during the summer (1981–2011, Fig. 3) while those in the Gulf of California consumed 88 primary prey species (1990–2019; Fig. 4) that varied between Zones (S2 Fig.). There was a lower number of prey species observed in the Channel Islands, despite the fact that diet diversity in this region using the Shannon Index is highest during summer (20). These regional differences in the number of prey items may mean that the ideal prey species were not consistently available for sea lions in the Gulf, or it may alternatively illustrate that there were more prey options available to facilitate diet adaptability in some Zones.

Overall, the Shannon Index and the total number of species consumed illustrate that the diet has been relatively consistent over time within the Channel Islands (Fig. 7, Table 1). In comparison, there was greater variability in diet composition in the Gulf of California both between Zones and over time within each Zone (S2 Fig.). For example, the number of species consumed per rookery ranged from 5–26 species between Zones (S5 Fig.), indicating greater resource heterogeneity and/or apparent dietary flexibility.

### 4.2 The role of diet energy density

Regardless of its relationship to diet diversity, the energy density of a diet is an important characteristic to consider when assessing diet quality. As each prey species differs in macronutrient composition and therefore in energy density (kJ/gww), diets based upon prey species that are more energy-rich can be considered “higher quality” as they are more likely to meet the nutritional requirements of individuals, allowing populations to grow. Measuring diet energy density can provide important insight into the nutritional status of a population and the drivers of population change.

On average, the diet energy density across the Gulf of California did not differ significantly from that of the Channel Islands, such that energy density did not explain broad differences in population trajectories. Contrary to expectations, we found that regions and periods when the diet had the highest energy density were not necessarily associated with years of greatest population growth within Zones. Alternate analyses (i.e., using finer data at the rookery-year scale and using changes in pup numbers as a more immediate indicator of changes in population demographics) also did not reveal any relationships between diet energy density and population changes. Even the diets with the highest energy densities were associated with both increasing and decreasing population trajectories, indicative of the lack of an overall simple relationship between diets and populations over time.

The highest mean diet energy density of all Zones (including the Channel Islands) occurred in the Gulf of California Zone 6 (San Esteban rookery), the largest rookery in terms of population size in the Gulf, where the sea lion population only showed a potentially increasing trend, while Zone 3 (Isla Lobos) had the second highest diet energy density but a significantly decreasing population. Furthermore, the only significantly increasing population in the Gulf of California (Zone 10, Los Islotes rookery) had a mean diet energy density that was comparable to the median diet energy density for all Zones (Table 1). While there is evidence from other pinniped studies to support the hypothesized link between diet energy density and population growth (24,40), it is possible that differences in diet quality in our study regions were not great enough to be the primary population drivers across the regions, except perhaps in cases with extremely low-quality diets (e.g., the decreasing population in Zone 4).

An important factor to consider when exploring the relationship between populations and diet quality is that, as demonstrated by our results, the Channel Islands and the Gulf of California are fundamentally different oceanographic systems with different population and diet dynamics. In the Channel Islands, most of the diet energy density consistently comes from schooling fishes (S1 Fig.), whereas there is considerable variability in the diet of sea lions in the Gulf of California, which is made up of different combinations of benthic species (41 different benthic species in total) making up their main source of energy from food (S2 Fig.), even though this prey category has a lower average energy density than schooling fishes. In fact, the greater predictability of the prey available to sea lions in the Channel Islands than those in the Gulf of California may be a more important contributor to population dynamics, rather than differences in prey energy density per se.

The heterogeneity in diet between rookeries throughout the Gulf of California raises questions about the specific trade-offs and foraging strategies among sea lions breeding at different rookeries. In the Galápagos Islands, individual foraging strategies of Galápagos sea lions (*Zalophus wollebaeki*) influence the coping abilities of their population with evidence that some foraging strategies may be more advantageous than others during environmental changes (42). In contrast to pelagic foragers, benthic foraging Galápagos sea lions appear to be less affected by increased water temperatures, despite consuming prey that are lower in energy density. Such environmentally dependent fitness trade-offs could also be at play in the Gulf of California sea lion populations, which should be explored through further research on individual foraging strategies.

### 4.3 The effects of environmental changes on diet quality

From 2013–2015, a large-scale phenomenon of increased sea surface temperatures known as “The Blob” was first documented in Alaska and then traveled south along the Eastern Pacific. In 2015– 2016, The Blob coincided with a strong El Niño event, further intensifying the effect of increased water temperatures (43). In the Channel Islands, El Niño events are known to cause population declines (with rapid subsequent recovery), and are associated with a decrease in consumption of energy-rich schooling fish (13). The Blob’s effects on the Channel Island populations were first seen in 2013 through increased pup mortality (20).

Unlike the Pacific coast, sea lions within the Gulf of California are not affected in the same way by warming events in the Pacific Ocean, such as El Niño. Instead, they appear to be affected by local oceanographic processes within the Gulf’s sub-regions (5). Overall, although the increased water temperatures in the Gulf after 2014 were characterized by an increase in the mean and range of diet diversity within the Gulf of California (expressed using the Shannon Index), this change was not significant (Fig. 8, top panel). However, there was an increase in the total number of species present in the diet (from 51 to 65 species, Table 2). In addition to the increase in the number of species consumed, the proportion of each diet species category changed (Fig. 5), as well as the identity of the species within those categories. For example, around 50% of the species consumed after 2014 were not present in the diet in previous years. These species consisted mostly of new benthic and lanternfish species (Figs. 7 and S6).

This increase in diet diversity was accompanied by a significant decrease in the overall average diet energy density throughout the Gulf of California (Fig. 8, bottom panel). This trend was driven by central rookeries (Zones 4 and 5) where there was a decrease in the proportion of energy-rich schooling fishes after 2014 (Fig. 9). Although there was also an increase after 2014 in the overall number of high-energy lanternfish species in the diet (from 1 to 6 species; Fig. 5), there was no significant change in their proportion (6% vs. 5%). This could suggest sea lions attempted to continue to meet their total energy requirements by increasing the diversity of energy-rich lanternfish species in their diet.

**Fig. 9.**
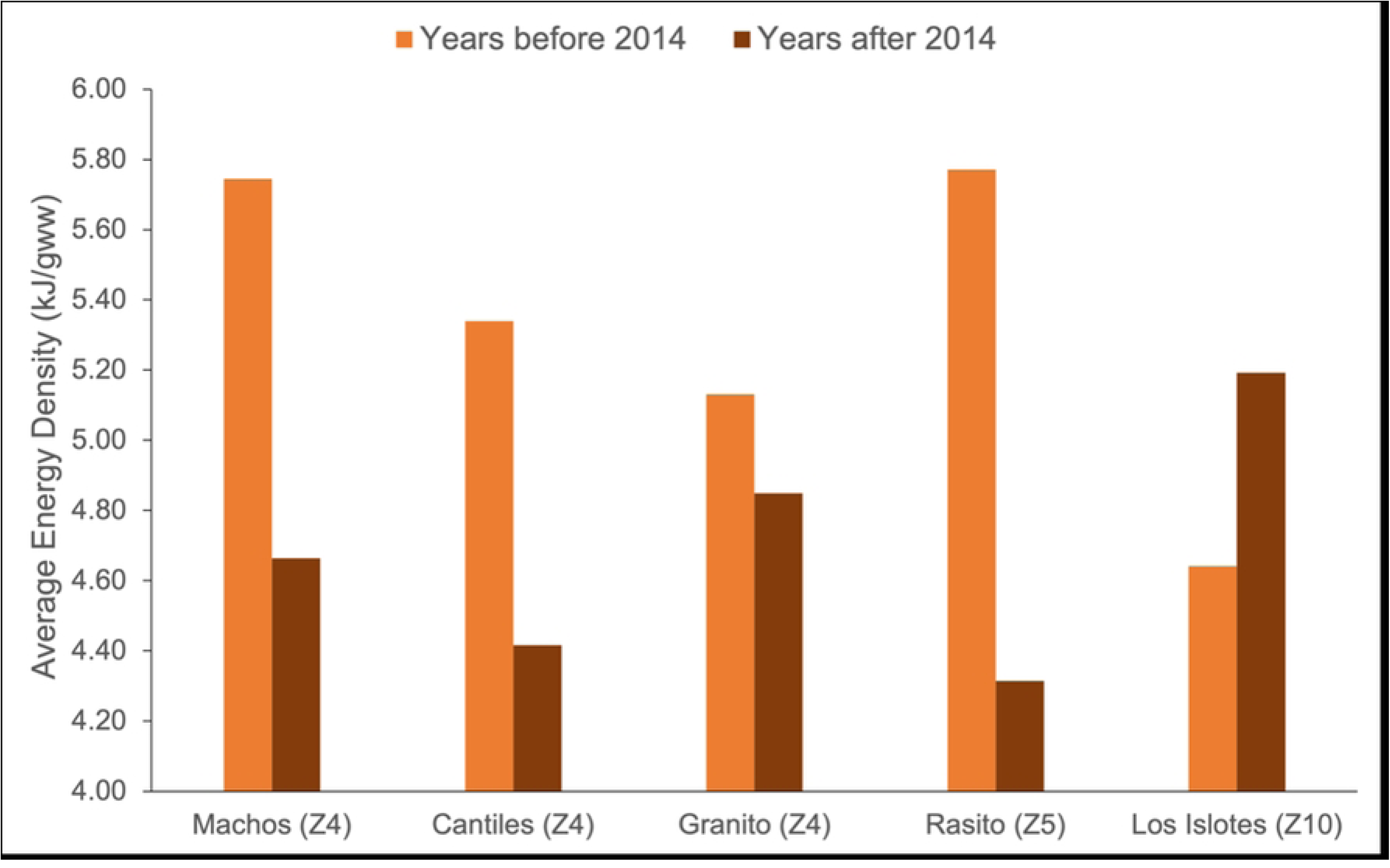
Average energy density of California sea lion diets at rookeries in the Gulf of California before and after 2014 from frequency of occurrence data. Only rookeries within Zones with matched data before and after 2014 are included in this Fig., their Zone number is shown in brackets. Gulf of California Zones are ordered geographically from North (Z4) to South (Z10). Zones that are not shown here lacked data after 2014 and were excluded from this Fig.

However, this pattern of decreased diet energy density after 2014 (mainly due to a decrease in energy-rich schooling fishes and lanternfish) was not consistent across all rookeries. A *decrease* in diet energy density after 2014 occurred at Rasito (Zone 5), which had a lack of energy-rich lanternfish and Jack mackerel in 2016 compared to 1996 (S2 Fig.). Conversely, within Zone 10 (Los Islotes rookery), an increase in lanternfish in 2019 was largely responsible for the overall *increase* in diet energy density after 2014 (Fig. 9).

The observed differences in the changes in diet energy density post-2014 among different Zones could be due to a difference in the availability of prey species in the various regions in the Gulf of California. In some Zones this could lead to California sea lions having to adopt atypical, lower-energy diets. For example, the unusual dominance of ‘other’ species in the diet in Los Islotes in 2015 may reflect a loss of ideal primary prey due to increased water temperatures (S2 Fig.), resulting in them consuming a higher number of prey species (S5 Fig.) to maintain the same level of dietary energy density from prey (∼5kJ/gww).

Previous studies have demonstrated how acute environmental changes and subsequent prey availability shifts can affect marine mammal population growth. For example, ringed seals (*Phoca hispida*) in western Hudson Bay switched to a more diverse diet that had a lower energy density due to decreases in the availability of their main prey (sand lance) triggered by changes in the seasonal breaking up of sea ice (40). As a result, body condition of individual seals was greatly reduced, and population declines ensued.

In the Channel Islands, models predict that every 1°C increase in surface temperature could decrease the population growth rate of California sea lions in the U.S. by 7% (44). Of note, during years of increased sea surface temperatures (2013–2015), (20) reported how sea lion diet composition in the Channel Islands decreased in epipelagic species (schooling fish), and increased in benthic and demersal species. A similar phenomenon appears to be occurring in the northern and central regions of the Gulf of California where California sea lion pup birth rates declined as sea surface temperature anomalies exceeded 1°C (17). However, population growth in the southern regions of the Gulf was not affected by increased sea surface temperatures (45), although pup abundance and body condition at Los Islotes did decrease during 2014 and 2015, and adult females were away from the rookery for longer periods than normal (46). Lactating females appeared to have had to forage further away from the rookery, which cost them more time and energy. Thus, warmer sea surface temperatures affected both the diet and foraging behaviour of female California sea lions in the south (Los Islotes, Zone 10), which may have affected pups during the lactation period.

### 4.4 Environmental heterogeneity and its implications for species management

The variable diet quality and population trends detailed in our study suggest that sea lions at different rookeries, even those just within the Gulf of California, cannot be viewed nor managed as a homogeneous group. The Gulf of California is known to have considerable environmental heterogeneity (47–50), which may influence both the quality of sea lion diets as seen in our study and, ultimately, predator-prey dynamics at the rookery scale. It has been suggested, for example, that compared to the northern and central regions of the Gulf, the greater diversity of prey species present in the south may buffer rookeries like Los Islotes against detrimental environmental changes (50,51). Having access to a greater diversity of prey species would allow sea lions to compensate for prey that may no longer be available.

Prey availability and abundance in the Gulf of California varies by region and is not as consistent or as predictable as in the California Current System. Such variability may mask the ability to identify simple relationships between diet and population growth, such as those shown in other Eastern Pacific ecosystems like the Channel Islands or Alaska (23,32,37). This variability may also underly differences noted by others in terms of genetic differences between California sea lions, their foraging areas, and the oceanographic conditions they experience (4,17,34). Understanding diet and population dynamics in the Gulf of California may therefore require a more detailed understanding of sea lion prey dynamics, foraging strategies, and localized oceanographic changes.

The Mexican government deems the California sea lion sub-populations in need of special protection, and recognizes the need to recover and conserve the populations in Mexico’s rookeries (52). Our study highlights the variation in the diets, population trajectories, rookery sizes, and oceanographic dynamics within the Gulf of California, suggesting that each rookery population faces different sets of challenges that impact their reproduction and survival rates in different ways.

However, current management and surveillance programs (especially in the central region) do not seem sufficient to monitor and assess how sea lion numbers are affected. More rigorous monitoring is needed not only to understand changes in prey species, but also other short-term factors affecting sea lion numbers such as entanglements in fishing gear, shootings, and contaminants to name a few (53). Such anthropogenic factors may also contribute to the lack of a direct relationship between diet and population trends in the northern and central Gulf of California (17). Overall, understanding the complex dynamics affecting each sub-population in both the short-term and long-term is essential to effectively manage the protection and conservation of California sea lions in the Gulf of California.

## Acknowledgements

The present work was part of Ana Lucía Pozas Franco’s Master’s thesis. Funding consisted of an NSERC Discovery Grant to Dr. David Rosen. Thank you to Dr. Miram Gleiber for sharing key energy density data unpublished at the time, Dr. Tony Orr for initial input and guidance, Eric Lee for help categorizing prey species, and Pedro González Espinosa for help in creating the Zone map.

## Supplementary Methods

### Calculating population change rules

In instances where a rookery had diet data for 3 or more continuous years, population data from one year before to one year after the diet data years were incorporated into the population change calculation. If the number of continuous years with diet data was less than three, then the population change was calculated from two years before and after the interval (or single year) of diet data. This yielded a single rate of population change over the matching diet data interval. In cases where population changes using the previously mentioned rules were calculated to be greater than ±20% (an unrealistic growth rate under normal breeding conditions), four years on either side of the diet data were incorporated into the calculation to obtain a more realistic rate of population change.

### Estimating missing population totals

Rookeries and years with available diet data were paired with available population totals. In some cases, population totals had to be estimated for years that lacked data. For the Channel Island rookeries, pup counts were available for years with missing population totals, and were therefore used to estimate totals by extrapolating from the linear relationship between pup counts and total population counts across all years. For the Gulf of California rookeries, missing population numbers were estimated by extrapolating from a linear regression performed on all available population data for individual rookeries because additional years with pup count data were not available. The regression equation was then used to estimate population numbers for years lacking counts, and the rate of population change over the period of interest was calculated from this mixed data set. Inaccurate extrapolations were avoided by only using regressions that spanned intervals with actual counts.

In cases where rookeries had one year of diet data and where the associated population data range (when incorporating the ±2 years rule for population change) overlapped with the population data range for another year of diet data, *and* one of those diet data years did not have +2 years of data after (due to it being the latest population year with data), those years of diet data were grouped. For example, diet data from the Granito rookery from 2016 and 2018 were combined into one grouping, and the years 2014–2018 were used to calculate population change since 2018 was the latest year with population data.

### Calculating rookery-year groupings

We matched the available diet data at the rookery level with a rate of population change value calculated to correspond to the specific year or group of consecutive years with available diet data. Ideally, continuous diet data for all years and rookeries would have been available, and matching data groupings would have been strategically chosen. However, diet data were patchy in terms of both years and rookeries. As a result, sequential data points were compiled from each rookery to form specific ‘year-rookery groupings’ and were treated as independent data points used in all subsequent analyses (Table S2).

Most rookeries in the Channel Islands had diet data available over several consecutive years, which were grouped according to the continuity of the data (e.g., San Miguel 2009–2011). Diet data from several rookeries in the Channel Islands were already averaged over several years (Table S2). In these instances, those year-rookery groupings were kept and used when calculating corresponding population changes. Within the Gulf of California, most years with diet data were single isolated years that varied by rookery (e.g., Los Islotes 1990, 2000, 2015 and 2019; Rasito 1996 and 2016, etc.). In cases where two consecutive years of diet data were available, we averaged the diet data and grouped them to form one rookery-year grouping (e.g., San Esteban 1995–1996). In cases where non-continuous years with diet data were close in time such that their population change calculations overlapped, we also combined them to form a single rookery-year grouping (e.g., Los Islotes 2015, 2019).

**Figure S1.**
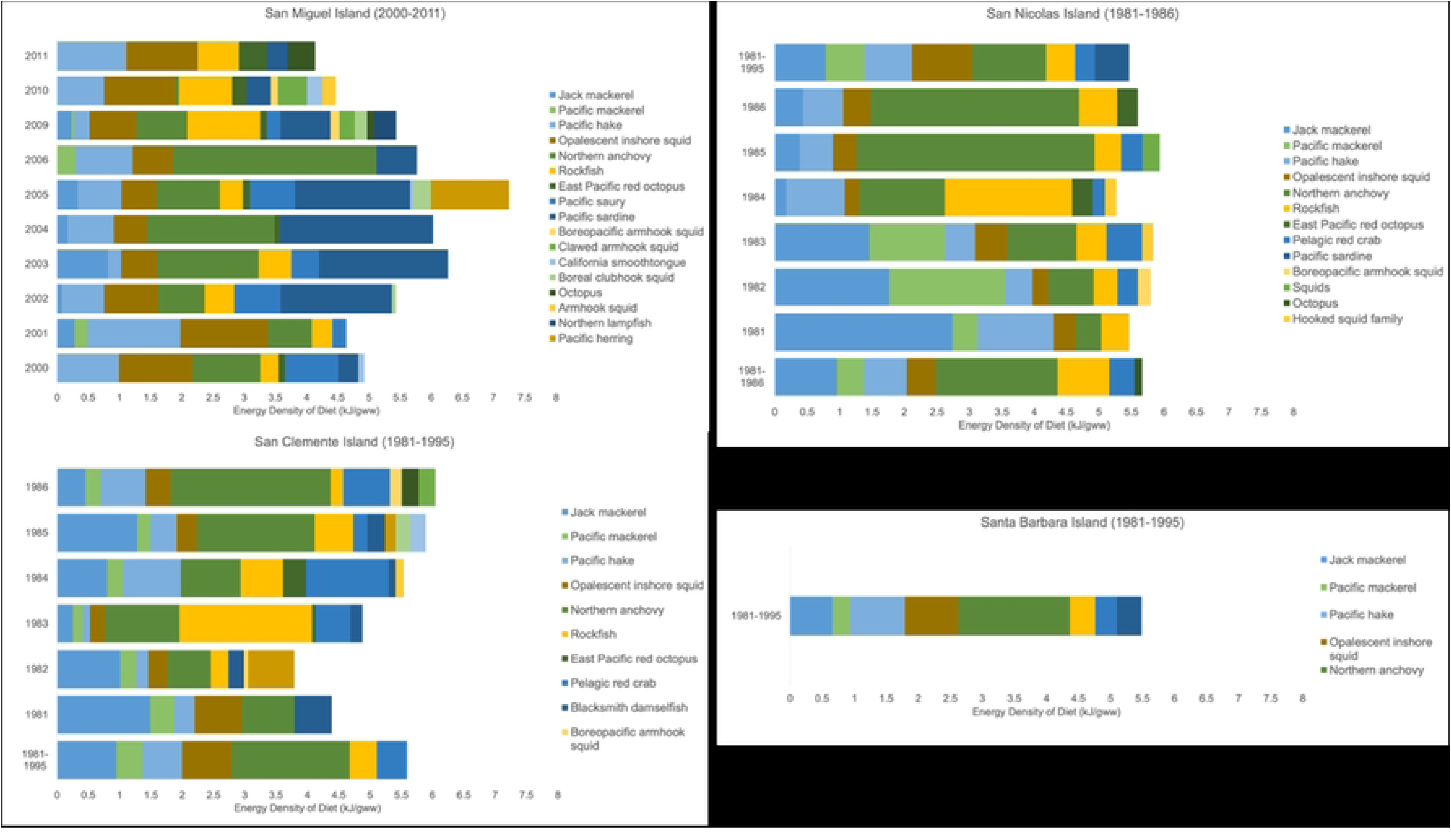
Energy density of diets of California sea lions in the Channel Islands (Zone 1). Average energy density and energetic content contributions (average weighted energy density) of the top 17 prey species to the total energetic content of the diet for rookeries and years with available frequency of occurrence data in the Channel Islands. ‘Other’ category represents all other species in the diet beyond the top 17.

**Figure S2.**
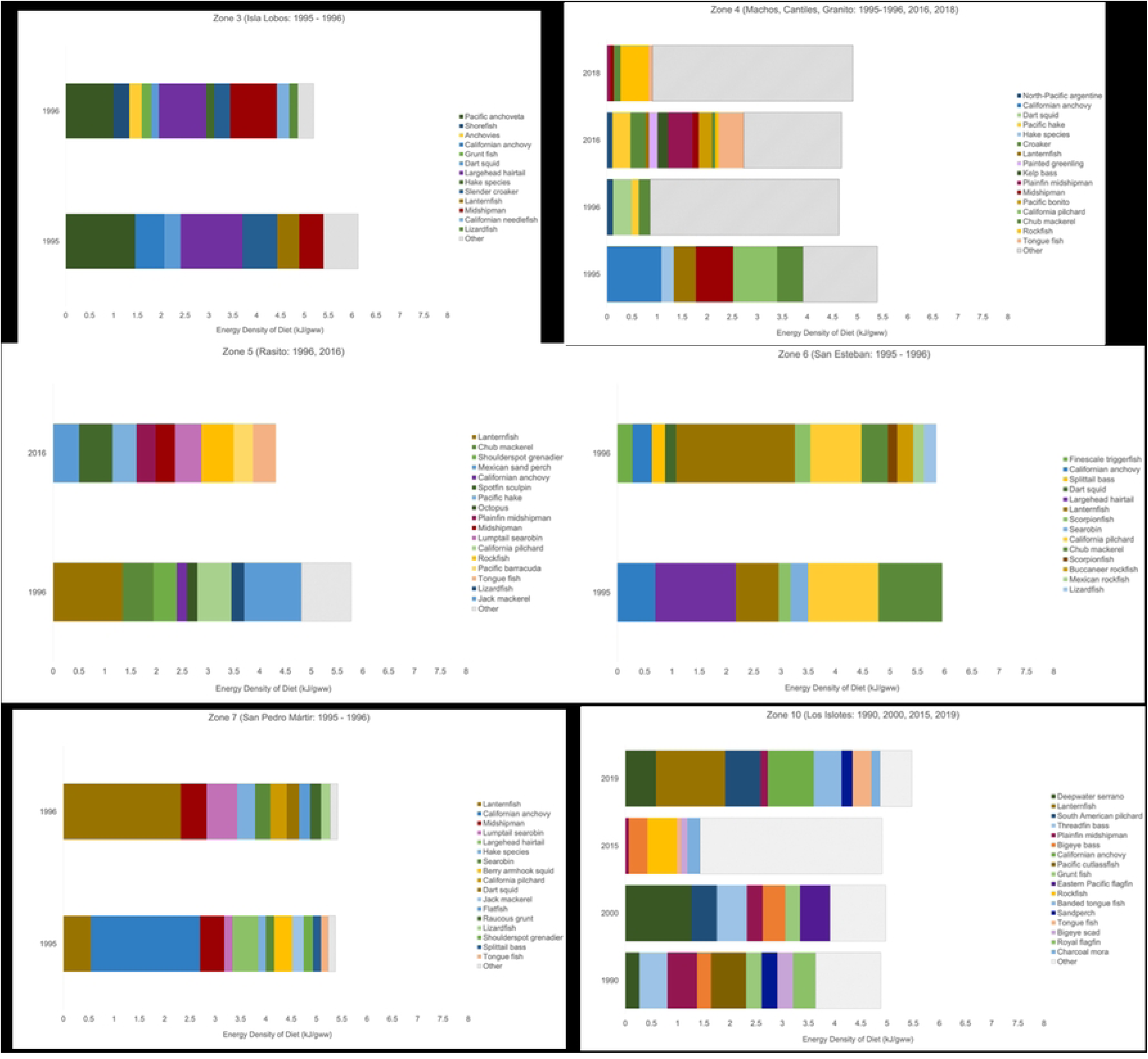
Energy density of diets of California sea lions in the Gulf of California (Zones 3– 5). Average energy density and energetic content contributions (average weighted energy density) of the top 17 prey species to the total energetic content of the diet for rookeries and years with available frequency of occurrence data in the Gulf of California. ‘Other’ category represents all other species in the diet beyond the top 17.

**Figure S3.**
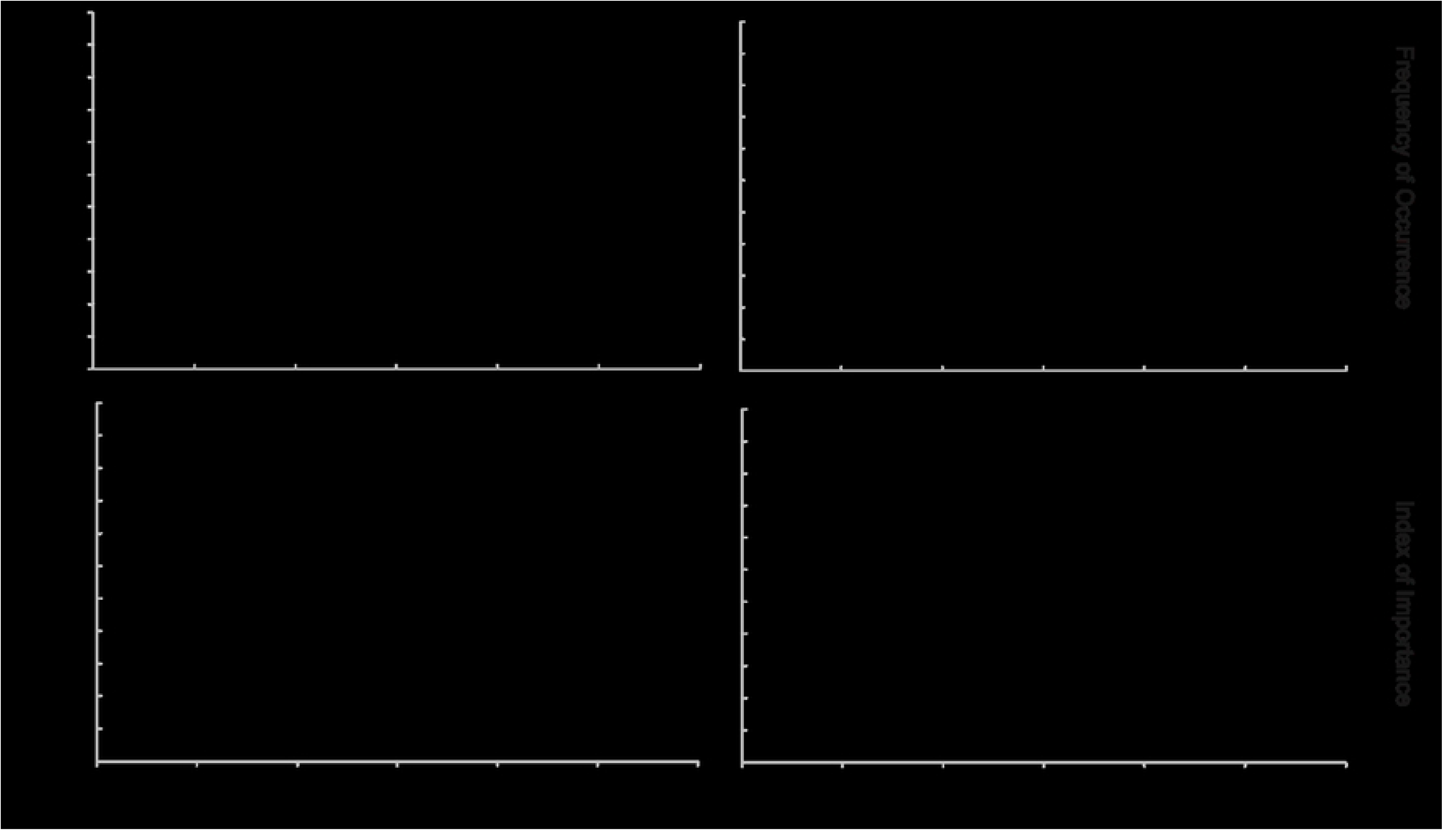
Population changes and diet quality for California sea lions from frequency of occurrence and index of importance data. The data represents values from Zone-era groupings. Diet diversity values were calculated using the Shannon Index from frequency of occurrence data (top-left panel) and index of importance data (bottom-left panel). Panels on the right include diet energy density values calculated using frequency of occurrence (top) and index of importance data (bottom). Regression analysis of data weighted by rookery population size indicated no statistical relationships.

**Figure S4.**
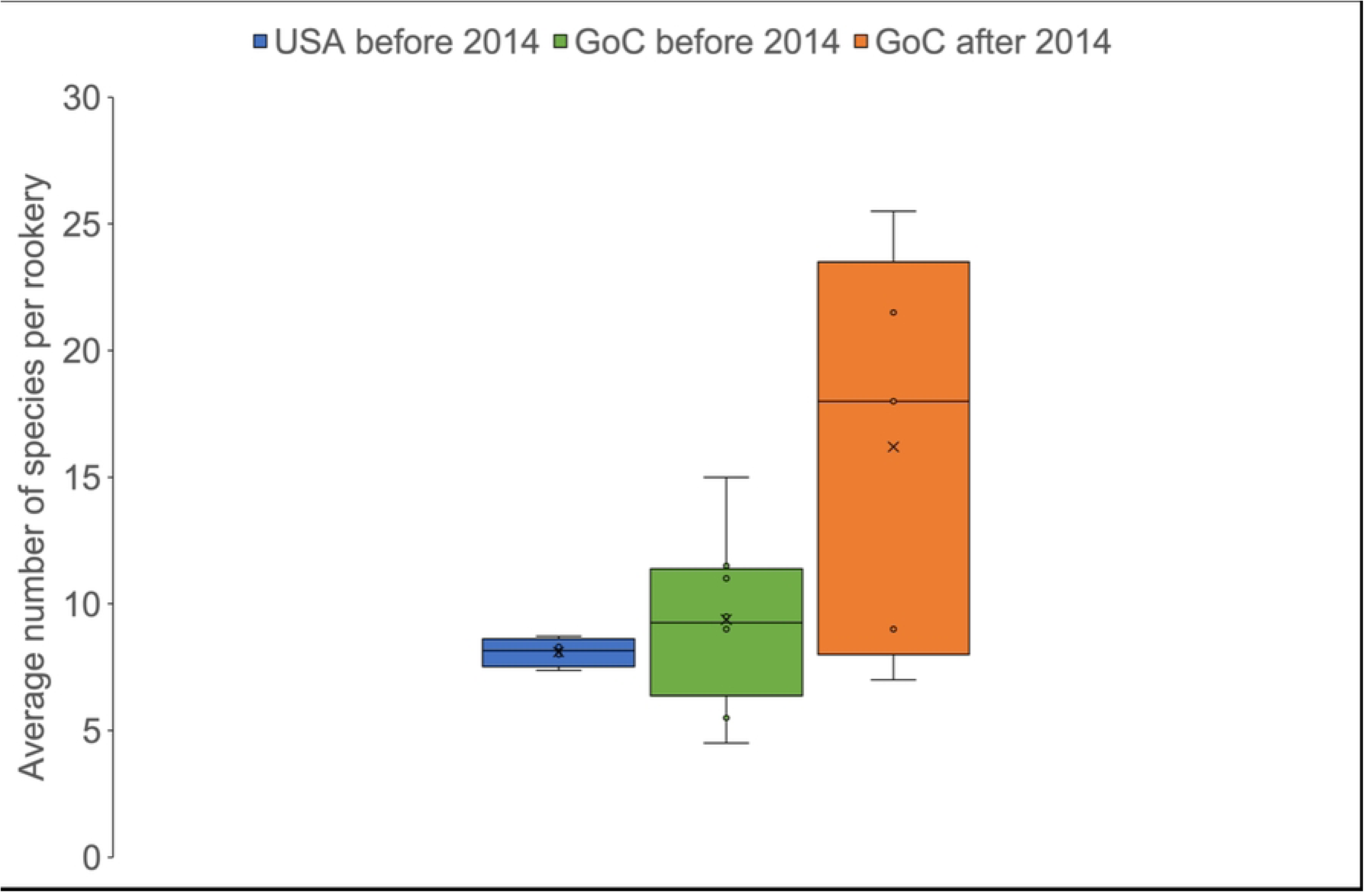
Average number of prey species per California sea lion rookery before and after 2014. Bars represent averaged data (mean value ± standard error). Average number of species is based on frequency of occurrence data. There was no data after 2014 available for the Channel Islands.

**Figure S5.**
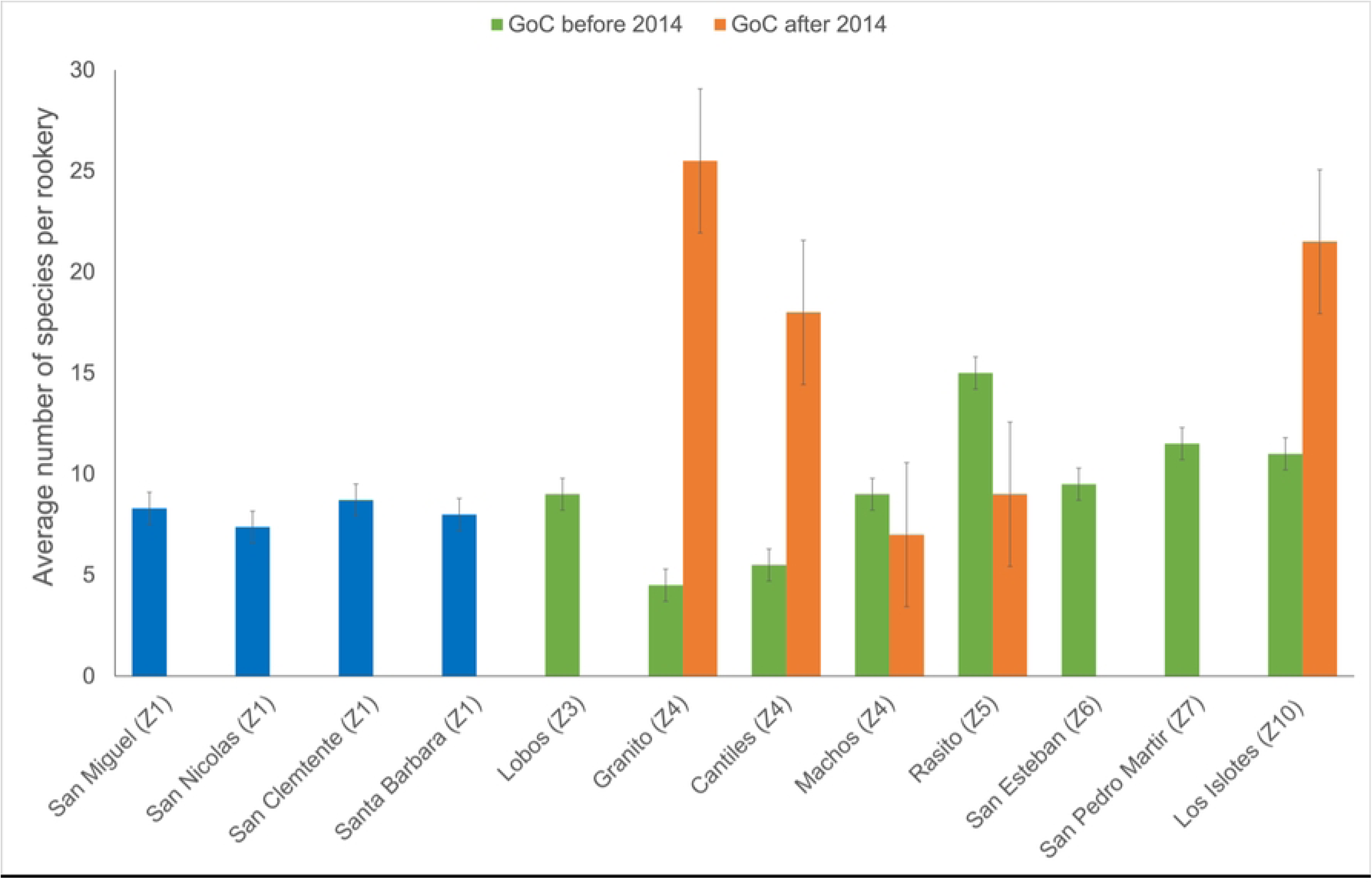
Average number of prey species per California sea lion rookery before and after 2014 from frequency of occurrence data. Blue bars represent each of the Channel Islands before 2014, each rookery with available data in the Gulf of California is shown before 2014 (green bars) and after 2014 (orange bars). Zone number is shown in brackets.

**Figure S6.**
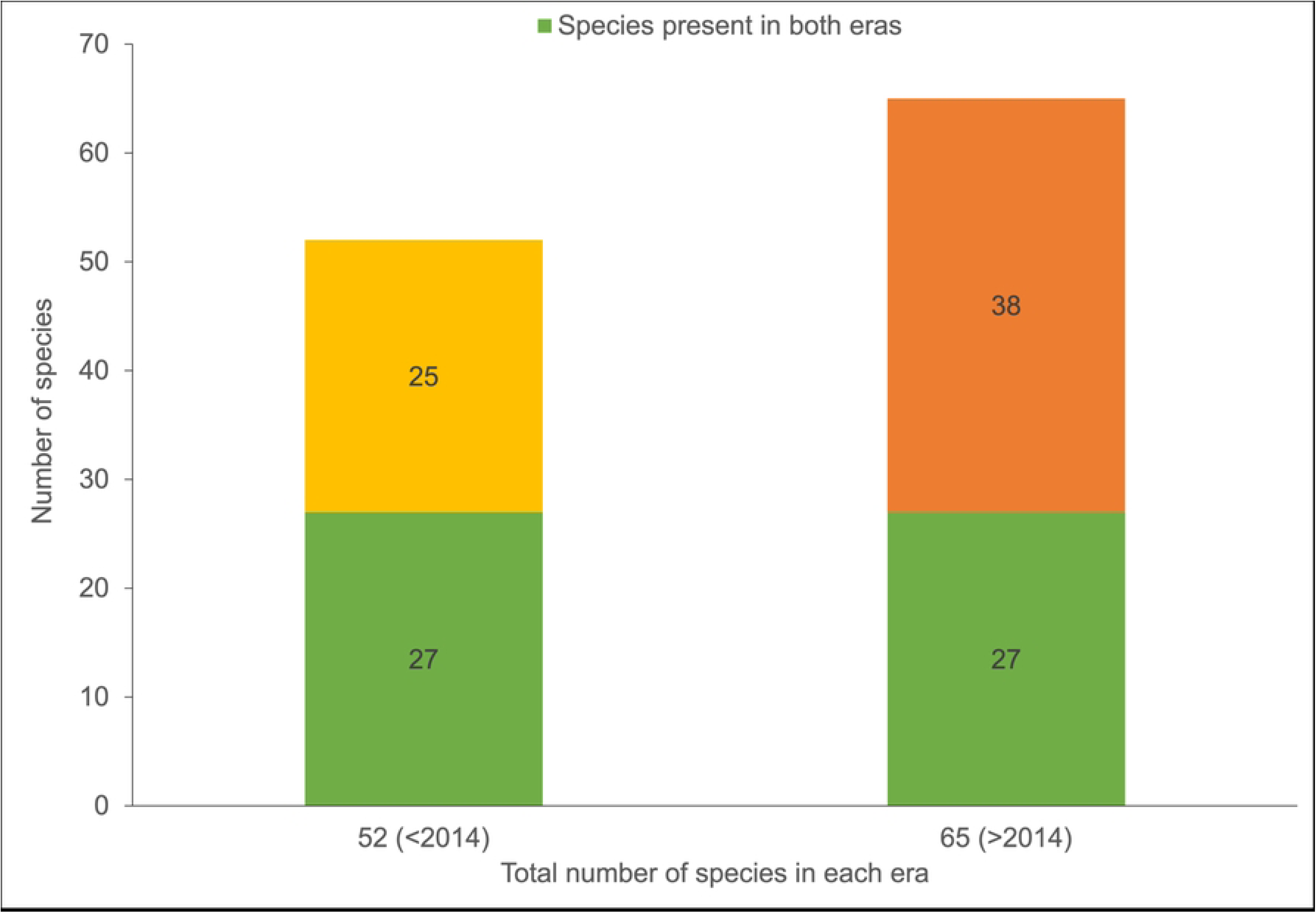
Total number of prey species consumed by California sea lions in the Gulf of California before and after 2014 from frequency of occurrence diet. The bar on the left represents prey species before 2014, the bar on the right represents prey species after 2014. Green bars show number of species present in both eras, yellow bar shows species only present before 2014, and orange bar shows number of species present only after 2014.

**Table S1.** Data sources of meta-analysis of California sea lion population and diet. Details include the rookery, data year and season, diet index or total population estimate (unless stated), and Table in source publication where relevant data is found. (FO; frequency of occurrence data, IIMP; index of importance data).

| Source | Rookeries | Year(s) of data | Season | Data type/diet index | Table |
| --- | --- | --- | --- | --- | --- |
| **Channel Islands** |  |  |  |  |  |
| Lowry et al. (2017a) | San Miguel | 1971–1991 (pup counts)<br>1992–2014 | July or August | Population (total live) | Table 2 |
|  | San Nicolas | 1991–2008<br>2009, 2010 (pup counts)<br>2011–2014 | July | Population (total live) | Table 2 |
|  | San Clemente | 1981–2014 | July or August | Population (total live) | Table 2 |
|  | Santa Barbara | 1983–1985 (pup counts)<br>1986–2008<br>2009, 2010 (pup counts)<br>2011–2014 | July | Population (total live) | Table 2 |
| Lowry et al. (1991) | San Nicolas | 1981–1986 (grouped) | June and August | FO | Table 1, 2 |
| Lowry and Carretta (1999) | San Nicolas, Santa Barbara, San Clemente | 1981–1995 | Multiple months, summer (Santa Barbara) | FO | Table 3 |
| A. Curtis unpubl. data | San Nicolas, San Clemente | 1981–1986 | Summer | FO | N/A |
| Lowry et al. (2017b) | Channel Islands | 2015 | July | Population (total counts) | Table 3 |
| Lowry et al. (2021) | Channel Islands | 2016–2019 | July or August | Population (total live) | Table 1 |
| Orr et al. (2011) | San Miguel | 2002–2006 | March–July | FO | Table 3 |
| S. R. Melin et al. (2012) | San Miguel | 2000–2003, 2005, 2009–2011 | June–Sept | FO | Table 5 |
| S. R. Melin et al. (2010) | San Miguel | 2000, 2001, 2002, 2004, 2005 and 2009 | July–early August | FO | Table 1 |
| **Gulf of California** |  |  |  |  |  |
| Pelayo-González, González-Rodríguez, et al. (2021) | 1–13 (not all rookeries have data for all years) | 1980–2019 | June or July | Population (total) | Raw data |
| García-Rodríguez (1995) | 13 | 1990 | February–September | FO | Table 4 |
| **Gulf of California** |  |  |  |  |  |
| F.J Garcia-Rodriguez unpubl. data | 5 | 1995 | September | FO & IIMP | N/A |
| F.J Garcia-Rodriguez unpubl. data,<br>Cardenas-Palomo (2003) | 3, 4, 6, 9, 10 | 1995 | June | FO & IIMP | Cuadro 1, Anexo 3 |
|  | 3, 4, 5, 8, 9, 10 | 1996 | May |  |  |
|  | 13 | 2000 | May 2000–April 2001 |  |  |
| Porras-Peters (2004) and Porras-Peters et al. (2008) | 1, 3, 5, 7–13 | 2002 | Summer | IIMP | Figure 4 (Appendix II) |
| Pelayo-González et al., (2021) | 13 | 2015 | July | FO | Raw data |
|  | 4 | 2016 | October |  |  |
|  | 5 | 2016 | Unknown |  |  |
|  | 6 | 2016 | October | IIMP |  |
|  | 8 | 2016 | October | FO & IIMP |  |
|  | 4 | 2018 | August | FO |  |
|  | 4 | 2018 | July |  |  |
|  | 5 | 2018 | July |  |  |
|  | 13 | 2019 | August |  |  |

Gulf of California rookeries: 1: Rocas Consagradas, 2: San Jorge (no data), 3: Isla Lobos, 4: Granito, 5: Cantiles, 6: Machos, 7: El Partido, 8: El Rasito, 9: San Esteban, 10: San Pedro Mártir, 11: San Pedro Nolasco, 12: Farallón de San Ignacio, 13: Los Islotes.

**Table S2.** Raw paired diet and population data for California sea lions by rookery-year grouping.

| **Rookery-year groupings** | **Zone** | **Index** | **Average diet diversity** | **Weighted energy density (kJ/gww)** | **Population change** | **Average energy density (kJ/gww)** | **Pup change (%)** |
| --- | --- | --- | --- | --- | --- | --- | --- |
| San Miguel 2000–2006 | 1 | FO | 1.70 | 5.76 | 5.03% | 5.6 | 3.2% |
| San Miguel 2009–2011 | 1 | FO | 2.00 | 4.68 | 7.80% | 5.1 | 11.6% |
| San Nicolas 1981–1986 (grouped) | 1 | FO | 1.92 | 5.66 | -2.92% | 5.36 | -4.8% |
| San Nicolas 1981–1986 | 1 | FO | 1.68 | 5.65 | -2.92% | 5.36 | -4.8% |
| San Nicolas 1981–1995 | 1 | FO | 1.95 | 5.46 | 9.47% | 5.85 | 32.0% |
| Santa Barbara 1981–1995 (grouped) | 1 | FO | 1.87 | 5.48 | 9.95% | 5.85 | 24.8% |
| San Clemente 1981–1995 (grouped) | 1 | FO | 1.81 | 5.59 | 4.51% | 5.74 | 7.7% |
| San Clemente 1981–1986 | 1 | FO | 1.96 | 5.09 | -1.61% | 5.50 | -1.9% |
| Los Islotes 1990 | 10 | FO | 2.40 | 4.59 | 2.43% | 4.7 | 5.12% |
| San Pedro Mártir 1995–96 | 7 | FO | 2.14 | 5.3 | -1.53% | 4.93 | -0.18% |
| San Esteban 1995–96 | 6 | FO | 2.04 | 5.9 | 7.29% | 5.08 | -1.52% |
| Rasito 1996 | 5 | FO | 2.48 | 5.77 | -7.60% | 5.15 | 2.31% |
| Machos 1995 | 4 | FO | 1.96 | 5.74 | -1.17% | 5.29 | 4.37% |
| Cantiles 1995–96 | 4 | FO | 1.36 | 4.51 | 6.15% | 5.05 | 6.76% |
| Granito 1995–96 | 4 | FO | 0.90 | 5.34 | 3.70% | 5.15 | 12.74% |
| Isla Lobos 1995–96 | 3 | FO | 1.93 | 5.13 | -2.03% | 5.57 | 5.16% |
| Los Islotes 2000 | 10 | FO | 2.03 | 4.69 | 7.37% | 5.00 | 3.26% |
| Rasito 2016 | 5 | FO | 2.20 | 4.31 | 7.25% | 4.31 | 13.52% |
| Machos 2016 | 4 | FO | 2.01 | 4.66 | -11.65% | 4.92 | 17.32% |
| Cantiles 2016 | 4 | FO | 2.70 | 4.42 | 7.09% | 4.58 | 7.92% |
| Granito 2016, 2018 | 4 | FO | 3.02 | 4.85 | 7.09% | 5.15 | 12.71% |
| Los Islotes 2015, 2019 | 10 | FO | 1.59 | 5.19 | -1.18% | 5.1 | 8.44% |
| San Pedro Mártir 1995–96 | 7 | IIMP | 1.526 | 5.86 | -1.5% | 5.58 | -0.2% |
| San Esteban 1995–96 | 6 | IIMP | 1.315 | 6.29 | 7.3% | 6.48 | -1.5% |
| Rasito 1995–96 | 5 | IIMP | 1.682 | 5.19 | -11.2% | 5.50 | 3.9% |
| Machos 1995 | 4 | IIMP | 1.528 | 5.94 | -1.2% | 5.75 | 4.4% |
| Cantiles 1995–96 | 4 | IIMP | 0.931 | 4.27 | 6.1% | 4.85 | 6.8% |
| Granito 1995–96 | 4 | IIMP | 0.561 | 5.30 | 3.7% | 5.37 | 12.7% |
| Isla Lobos 1995–96 | 3 | IIMP | 1.479 | 5.19 | -2.0% | 4.47 | 5.2% |
| Los Islotes 2002 | 10 | IIMP | 1.848 | 4.7 | 4.4% | 5.08 | -0.2% |
| San Esteban 2002 | 6 | IIMP | 1.503 | 5.3 | 9.2% | 5.42 | 16.3% |
| Rasito 2002 | 5 | IIMP | 0.062 | 5.43 | 5.5% | 4.85 | 12.9% |
| San Pedro Mártir 2002 | 7 | IIMP | 0.685 | 3.10 | -0.7% | 4.98 | -6.4% |
| Cantiles 2002 | 4 | IIMP | 1.096 | 3.81 | -3.6% | 3.81 | -8.8% |
| Isla Lobos 2002 | 3 | IIMP | 1.326 | 4.71 | 4.5% | 5.01 | -2.1% |
| San Pedro Nolasco 2002 | 8 | IIMP | 1.874 | 4.99 | -1.4% | 5.11 | -2.1% |
| Partido 2002 | 5 | IIMP | 1.454 | 6.81 | -7.9% | 5.29 | -3.9% |
| Rocas Consagradas 2002 | 2 | IIMP | 1.060 | 5.33 | 5.4% | 4.63 | 5.4% |
| Farallón de San Ignacio 2002 | 9 | IIMP | 2.386 | 6.20 | -3.8% | 4.94 | -5.2% |
| Los Islotes 2015 | 10 | IIMP | 0.957 | 4.38 | 1.3% | 5.03 | 8.2% |
| Rasito 2016 | 5 | IIMP | 1.921 | 4.17 | 7.2% | 4.31 | 13.5% |
| Granito 2016, 2018 | 4 | IIMP | 1.300 | 4.59 | 6.6% | 4.84 | 12.7% |
| Cantiles 2016, 2018 | 4 | IIMP | 1.433 | 5.15 | 7.1% | 4.90 | 7.9% |
| Los Islotes 2019 | 10 | IIMP | 1.684 | 6.10 | -3.7% | 5.09 | 8.6% |

**Table S3.** Raw paired diet and population data for California sea lions by Zone-era grouping.

| **Zone-era groupings** | **Era** | **Index** | **Average diet diversity** | **Population change** | **Weighted average energy density (kJ/gww)** | **Median population** |
| --- | --- | --- | --- | --- | --- | --- |
| Zone 1 – Channel Islands | 1981–1995 | FO | 1.87 | 2.7% | 5.49 | 9,216 |
| Zone 1 – Channel Islands | 2000–2011 | FO | 1.85 | 6.4% | 5.22 | 44,720 |
| Zone 3 – Isla Lobos | 1995–1996 | FO | 1.93 | -2.0% | 5.13 | 2,822 |
| Zone 4 – Machos, Cantiles, Granito | 1995–1996 | FO | 1.41 | 2.9% | 5.20 | 1,355 |
| Zone 4 – Machos, Cantiles, Granito | 2016, 2018 | FO | 2.58 | 0.8% | 4.64 | 696 |
| Zone 5 – Rasito | 1996 | FO | 2.48 | -7.6% | 5.77 | 362 |
| Zone 5 – Rasito | 2016 | FO | 2.20 | 7.2% | 4.31 | 308 |
| Zone 6 – San Esteban | 1995–1996 | FO | 2.04 | 7.3% | 5.90 | 7,171 |
| Zone 7 – San Pedro Mártir | 1995–1996 | FO | 2.14 | -1.5% | 5.32 | 1,963 |
| Zone 10 – Los Islotes | 1990 | FO | 2.21 | 4.9% | 4.64 | 347 |
| Zone 10 – Los Islotes | 2015, 2019 | FO | 1.59 | -1.2% | 5.19 | 538 |
| Zone 2 – Rocas Consagradas | 2002 | IIMP | 1.06 | 5% | 5.33 | 839 |
| Zone 3 – Isla Lobos | 1995–1996 | IIMP | 1.48 | -2% | 5.19 | 2,822 |
| Zone 3 – Isla Lobos | 2002 | IIMP | 1.33 | 4% | 4.71 | 1,897 |
| Zone 4 – Machos, Cantiles, Granito | 1995–1996 | IIMP | 1.01 | 3% | 5.17 | 1,355 |
| Zone 4 – Machos, Cantiles, Granito | 2002 | IIMP | 1.10 | -4% | 3.81 | 1,090 |
| Zone 4 – Machos, Cantiles, Granito | 2016, 2018 | IIMP | 1.20 | 7% | 4.87 | 729 |
| Zone 5 – Rasito, Partido | 1995–1996 | IIMP | 1.68 | -11% | 5.19 | 366 |
| Zone 5 – Rasito, Partido | 2002 | IIMP | 0.76 | -1% | 6.12 | 507 |
| Zone 5 – Rasito, Partido | 2016 | IIMP | 1.30 | 7% | 4.17 | 308 |
| Zone 6 – San Esteban | 1995–1996 | IIMP | 1.32 | 7% | 4.99 | 7,171 |
| Zone 6 – San Esteban | 2002 | IIMP | 1.45 | 9% | 6.81 | 6,334 |
| Zone 7 – San Pedro Mártir | 1995–1996 | IIMP | 1.53 | -2% | 5.86 | 7,171 |
| Zone 7 – San Pedro Mártir | 2002 | IIMP | 0.69 | -1% | 3.10 | 2,405 |
| Zone 8 – San Pedro Nolasco | 2002 | IIMP | 1.87 | -1% | 4.99 | 937 |
| Zone 9 – Farallón de San Ignacio | 2002 | IIMP | 2.39 | -4% | 6.20 | 643 |
| Zone 10 – Los Islotes | 2002 | IIMP | 1.85 | 4% | 4.72 | 404 |
| Zone 10 – Los Islotes | 2015 | IIMP | 1.92 | 1% | 4.38 | 538 |
| Zone 10 – Los Islotes | 2019 | IIMP | 1.68 | -4% | 6.10 | 659 |

**Table S4.**
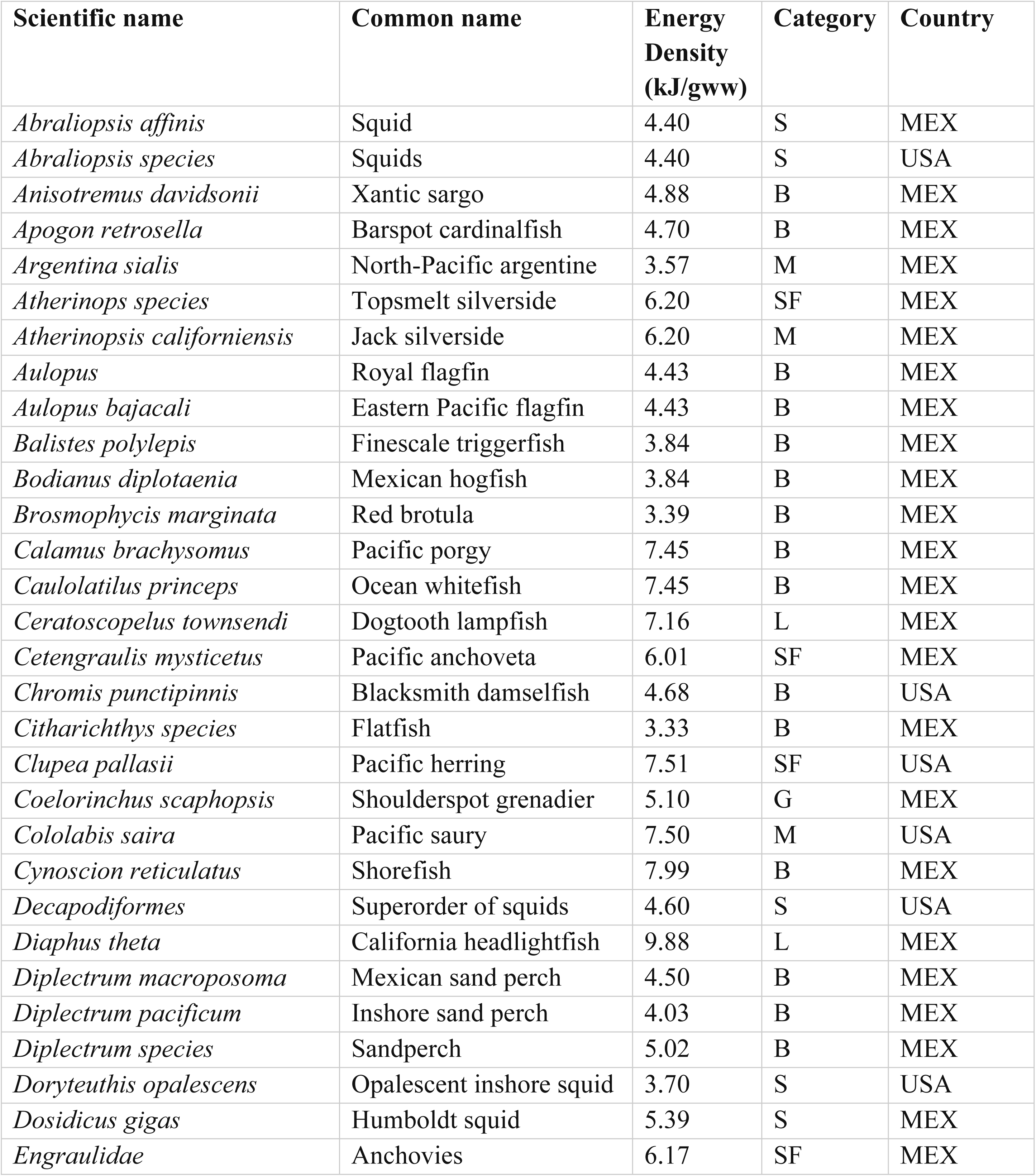
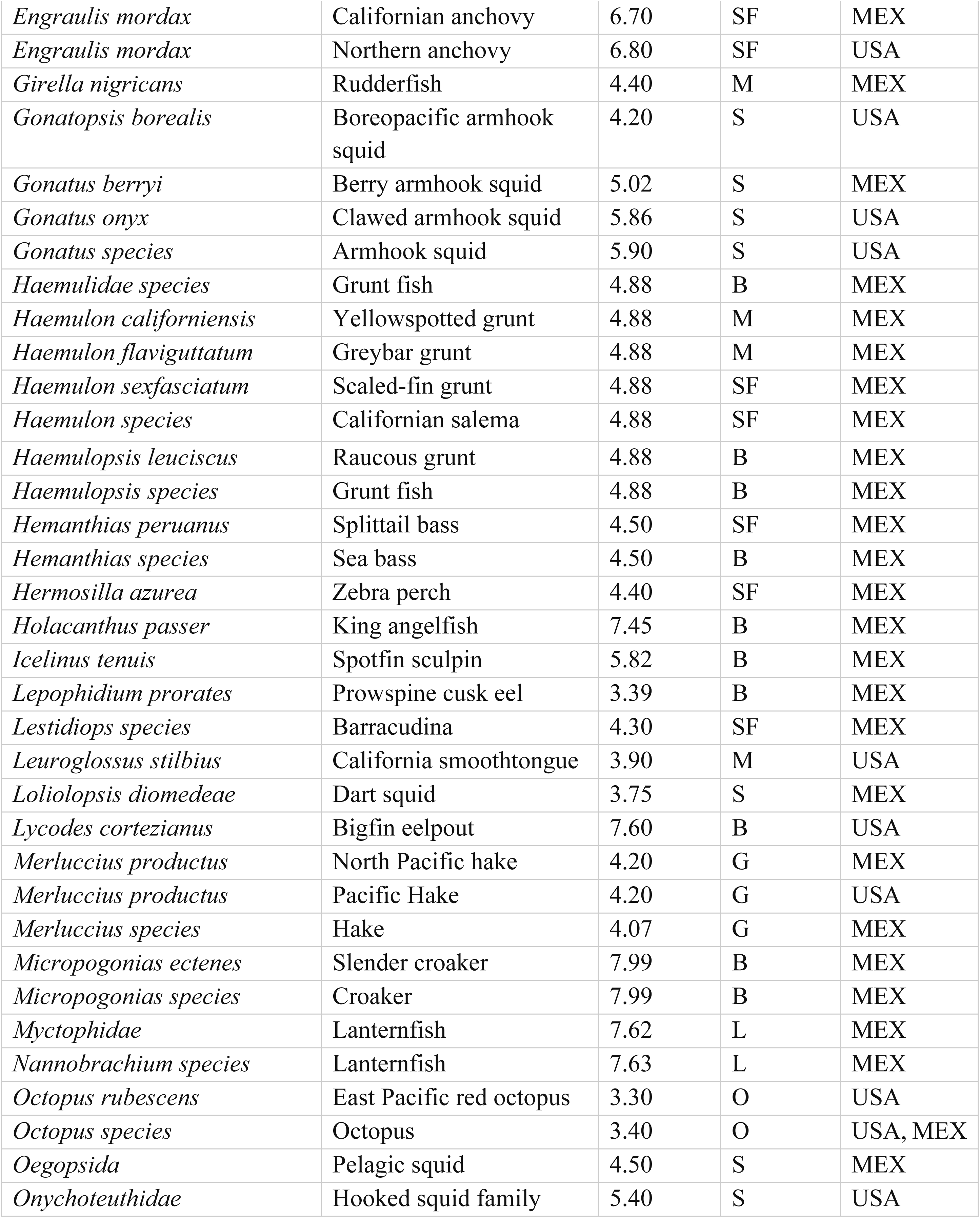
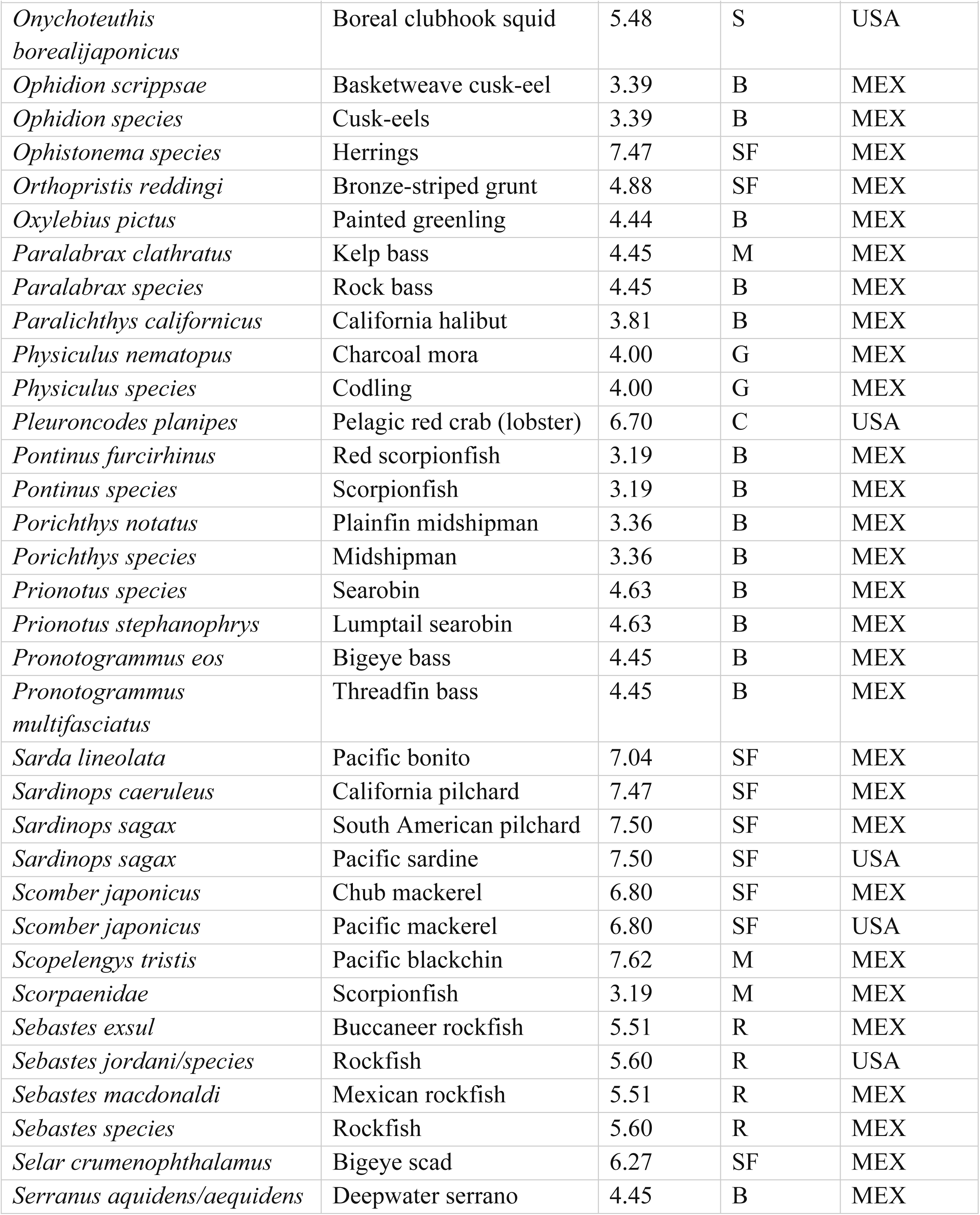
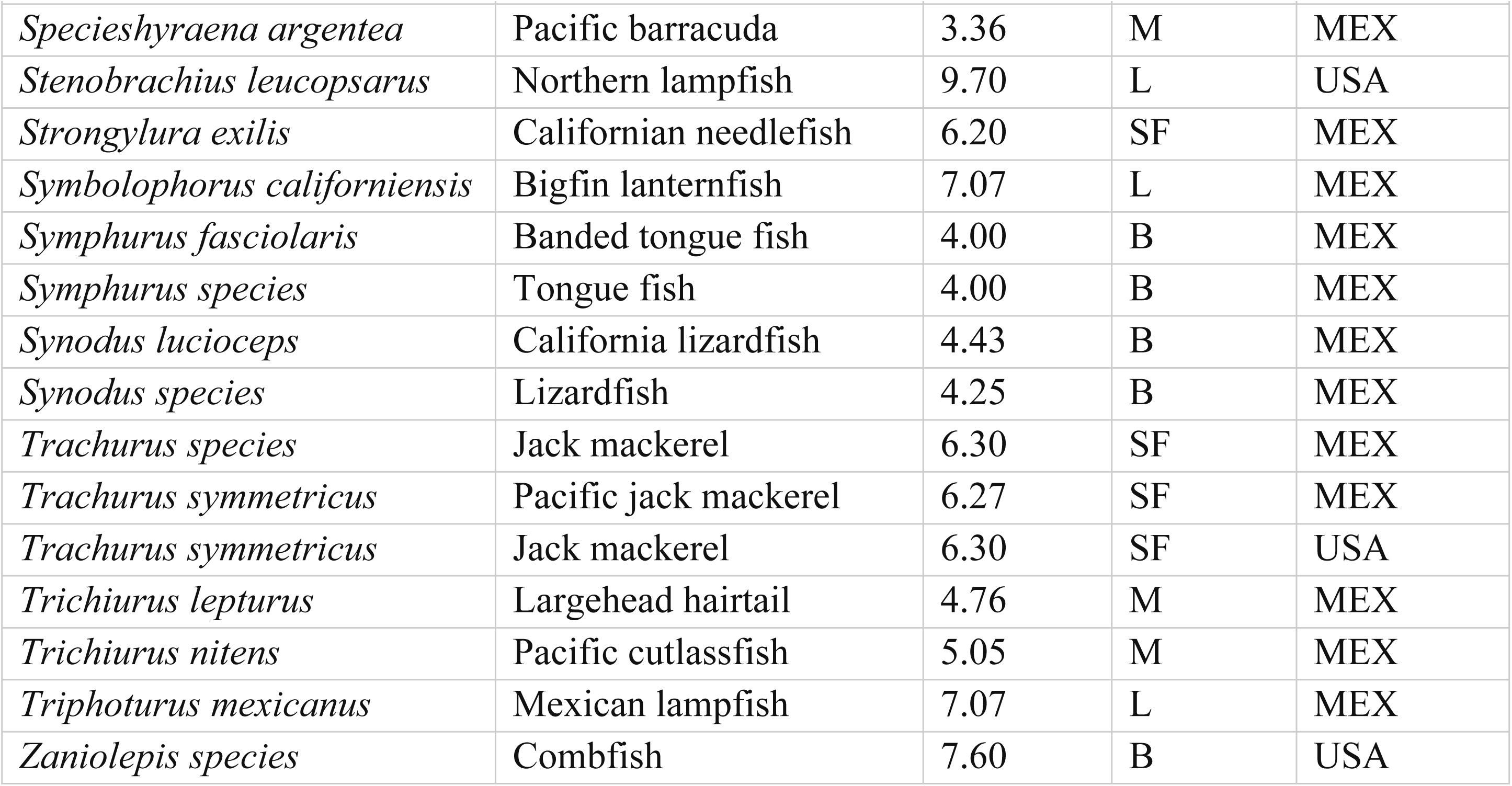
List of 114 diet prey species (scientific and common name) consumed by California sea lions in the Gulf of California (MEX) or the Channel Islands (USA) showing their average energy density, category assigned, and country where data was collected. Some species had the same scientific name, but different common names depending on the region. Categories: B: benthic species, C: crustaceans, G: gadids, L: lanternfish, O: octopus, R: rockfish, SF: schooling fish, S: squid, M: miscellaneous.

## Notes

### Competing Interest Statement

The authors have declared no competing interest.

